# Differential DNA methylation clock ages across buffy coat (BC), peripheral blood mononuclear cells (PBMC), and saliva in individuals in early-to-mid adulthood

**DOI:** 10.1101/2025.09.16.673560

**Authors:** Ryan Bruellman, Donald Evans, Andrew Smolen, Luke M. Evans, Chandra A. Reynolds

**Author notes:** Correspondence to: Chandra A. Reynolds, Institute for Behavioral Genetics, 1480 30^th^ Street, University of Colorado, Boulder, Colorado 80309, USA.

## Abstract

Epigenetic aging prior to midlife is gaining interest as an intervenable period to address health and cognitive aging. Epigenetic changes may index DNA methylation aging rates, but methylation profiles may not be substitutable across tissues. We compared DNA methylation clocks and age acceleration in saliva, buffy coat (BC), and peripheral blood mononuclear cells (PBMC) collected in 91 individuals (7 unpaired, 20 siblings, 64 twins; 18 monozygotic (MZ), 14 dizygotic (DZ) pairs) from the Colorado Adoption/Twin Study of Lifespan behavioral development and cognitive aging (CATSLife1; Mean age=30.90 years [range=28.07–41.13]; 50.5% female). Across 15 DNA methylation clocks, chronological age and DNA methylation ages were moderately associated (mean Spearman correlations: *r*=0.37, Saliva; *r*=0.41, BC; *r*=0.38, PBMC). In mixed-effects models, saliva showed higher DNA methylation ages (*B*=3.83–16.46 years vs BC, p<0.001), whereas PBMC and BC were comparable (*B*=-0.06–0.39 years vs BC, p≥0.447). The exception was the next-generation clock DunedinPace showing comparability (*p*=0.486). Similar patterns were observed for age acceleration estimates. Altogether, MZ pairs (meta-analytic *r=*0.49, 95%CI=0.32,0.66) and DZ pairs (meta-analytic *r=*0.38, 95%CI=0.25,0.52) were moderately correlated (Spearman), but MZ pairs showed heterogeneity across tissues (*p*<0.020): saliva was lower (mean *r=*0.33, *SD*=0.25) than BC (mean *r=*0.64, *SD*=0.10) and PBMC (mean *r=*0.49, *SD*=0.18). Next-generation PCGrimAge and DunedinPace clocks showed consistent zygosity correlations across tissues, while multi-tissue clocks (e.g., ZhangQ) showed comparable MZ-DZ correlations. While saliva-based DNA methylation is not a direct substitute for blood-based DNA methylation, BC and PBMC show comparability; nevertheless, all tissue types may be appropriate for DNA methylation aging studies when compared within tissues.

**Key Policy Highlights:**

- Saliva-based methylation is not a direct substitute for blood-based methylation.
- Saliva-based DNA methylation clocks demonstrated relatively older methylation ages and faster age accelerations relative to blood-based DNA methylation clocks.
- Blood-based DNA methylation clocks were comparable across buffy coat and peripheral blood mononuclear cell tissues.
- Twin similarity, particularly among identical twins, was higher for blood-based DNA methylation clocks and newer generation DNA methylation clocks.
- Three clocks (one first-generation multi-tissue and two next-generation) showed consistency and/or comparability across tissues and showed moderate to strong correlations among identical and fraternal twins, supporting the importance of tool development.

## Introduction

Individual differences in DNA methylation patterns may signal important variation in gene expression in response to exposures and may index accelerated aging. DNA methylation may increase or decrease gene expression over time (Singal & Ginder, 1999), where attachment of a 5-methyl group to cytosine-phosphate-guanine (CpG) sites leads to reductions or silencing of specific gene activity (Mattei, Bailly, & Meissner, 2022). The methylation process is a vital component of development and has wide implications across aging. While the DNA sequence remains relatively stable as we age (Schumacher, Pothof, Vijg, & Hoeijmakers, 2021), DNA methylation patterns differ by tissue type and are subject to alterations across time within tissue (Gopalakrishnan, Van Emburgh, & Robertson, 2008; Jones, Goodman, & Kobor, 2015; Miller et al., 2023; Robertson, 2005; Slieker, Relton, Gaunt, Slagboom, & Heijmans, 2018). Tissue selection in methylation research is often determined by ease of sampling, with blood and saliva most commonly used (Langie et al., 2017). Tissue selection is, nonetheless, a salient consideration that may illuminate possible mechanisms if proximally or directly associated with phenotypes of interest. Many studies rely on stored saliva or buffy coat (BC) samples as opposed to whole blood or peripheral blood mononuclear cells (PBMC), and such biobanked samples could be a boon for research into DNA methylation, especially where comparable (Dou et al., 2018; Ghamrawi et al., 2021; Langie et al., 2017; Staunstrup et al., 2017). Understanding whether and when significant differences in methylation measurements exist across the commonly collected biological mediums of saliva, BC, and PBMC can assist researchers to balance comparability with accessibility.

Because tissue-choice influences methylation profiles, researchers must account for the complexity this introduces when comparing sample types, even for those samples derived from the same parent tissue (e.g., blood in the case of BC and PBMC). Quality control in methylation analysis includes estimating cell counts to reduce inter-sample variability within tissue type (Teschendorff & Horvath, 2025). Nonetheless, cross-tissue differences remain pertinent due to distinct physiological roles. Saliva, commonly used for its ease in collection and low costs (e.g., Birknerova et al., 2022; Del Toro et al., 2024; Langie et al., 2017; Smith et al., 2015), is a composite tissue containing both epithelial cells and leukocytes (Theda et al., 2018). BC consists of PBMC components, including lymphocytes and monocytes, as well as granulocytes, platelets, and residual red blood cells (Martins et al., 2024; Navas, Giraldo-Parra, Prieto, Cabrera, & Gómez, 2019). PBMC is often the tissue of choice in DNA methylation work in studies of disease markers, disease progression, as well as aging (Cheishvili et al., 2024; Friso et al., 2013; Y. Li et al., 2010; Steegenga et al., 2014; Stefanowicz et al., 2012; T. Wang et al., 2023).

DNA methylation clocks are widely calculated across tissue types as biological aging indicators. The clocks differ as to the number of CpG sites measured, the locations of CpG sites, the chronological age ranges of the training samples, and the tissue type or types used to derive the clock (Y. Wang, Grant, Zhai, McDonald-Maier, & Schalkwyk, 2024). These clocks measure methylation activity across selected CpG sites to calculate a single summary value that may provide insights into biological aging and possible risks of aging-related disease and mortality (Jones et al., 2015). Clock metrics range from methylation age, the relative rate of aging, and mortality risk, to tracking biological process (e.g., telomere lengths, mitotic divisions). First-generation DNA methylation clocks have been trained on chronological age, using single or multiple tissues, while ‘next-generation’ clocks have been trained on age-related health phenotypes, producing methylation age or rate of aging estimators (Johnson & Shokhirev, 2025). Hence, chronological age is positively and strongly correlated with DNA methylation age in most cases (X. Li et al., 2020). Deviations from chronological age, either lower or higher, may indicate relatively slower or faster methylation aging and lesser or greater susceptibility to disease, respectively. Higher methylation age may signal a need for interventions to alter behavior, disease susceptibility, or disease progression (Gopalakrishnan et al., 2008; Robertson, 2005).

Despite their utility, questions remain about whether and when these DNA methylation clocks perform consistently across different tissue types. Early work suggested a similarity of DNA methylation profiles between saliva and blood tissues in most but not all studies (Langie et al., 2017; Staunstrup et al., 2017) with further testing needed. Recent comparison studies are small in sample size and focused on platform differences (Tay et al., 2025), employ only two tissue types (e.g., whole blood, saliva) (Zarandooz & Raffington, 2025), and/or compare multiple tissues across a wide age range (Apsley et al., 2025). For example, comparisons of buccal epithelial cells, saliva, dried blood spots (DBS), BC, and PBMC samples (Apsley et al., 2025), suggested salient differences between the three blood versus two oral tissue types in terms of DNA methylation clocks. However, sample sizes per tissue type varied between 35 - 81 individuals and ages ranged from 9 to 70 years of age, with BC available in children only (from a high-risk maltreatment sample) and PBMC in adults only. Of seven DNA methylation clocks, the pan-tissue Skin and Blood Clock (Horvath et al., 2018) exhibited the greatest concordance across all tissue types, with no differences in methylation ages in saliva-BC or saliva-DBS comparisons, but with older methylation ages for saliva versus PBMC (in the adult subsample only, N=45) (Apsley et al., 2025).

To address gaps, this cross-sectional study evaluates the consistency and comparability of multiple DNA methylation clocks across saliva, BC, and PBMC tissues, collected simultaneously in a genetically informative sample in established adulthood (28-41 years) reflecting early-to-mid-adulthood (c.f.,Mehta, Arnett, Palmer, & Nelson, 2020). The Colorado Adoption/Twin Study of Lifespan behavioral development and cognitive aging (CATSLife), is a genetically and environmentally informative study, including both monozygotic and dizygotic twins, as well as biological and adopted sibling pairs, allowing the dissection of genetic and environmental factors that contribute to age-related outcomes. The goal of the present study is to determine the comparability of tissue source in a cross-sectional sample to inform future DNA methylation studies. Specifically, we test differences among 15 DNA methylation clocks in BC, PBMC, and saliva in 91 individuals. Our sample includes a subset of 64 twins (32 pairs) which allows us to evaluate the relative similarity of monozygotic (MZ) to dizygotic (DZ) twin pairs. We predicted that BC and PBMC based clocks will more closely align with each other than with saliva based clocks. Moreover, we expected that first-generation clocks trained on multiple tissues or next-generation clocks trained on multiple aging related biomarkers (Belsky et al., 2022; Horvath et al., 2018; Lu et al., 2019; Zhang et al., 2019), would show stronger associations across sample mediums (Apsley et al., 2025; Zarandooz & Raffington, 2025). Last, we expected that MZ twins would show greater similarity than DZ twins (Jylhava et al., 2019; Reynolds et al., 2020), with greater twin similarity for more recently developed clocks.

## Methods

### Participants

Samples from 91 participants were selected as a subset of the Colorado Adoption/Twin Study of Lifespan behavioral development and cognitive aging (CATSLife1) conducted between 2015 to 2021 (Wadsworth et al., 2019). The study has adhered to the Declaration of Helsinki guidelines and was approved by the Institutional Review Boards at the at the University of Colorado Boulder [14-0421] and the University of California, Riverside [HS 14-073]. Written informed consent was obtained from each participant. Among these 91 participants, 84 individuals were from sibling or twin pairs including 20 sibling pairs (seven biological pairs, three adoptive pairs), 14 dizygotic (DZ) twin pairs, and 18 monozygotic (MZ) twin pairs. The sample was roughly evenly split by sex (46 female, 45 male) with 80 participants self-reporting their race as White and 11 as Multiple/other, and 80 participants self-reporting their ethnicity as non-Hispanic and 11 as Hispanic.

Participant chronological age at the time of blood draw ranged from 28.07 - 41.13 years with an average age of 30.90 years (*SD* = 3.75 years). Differences in chronological ages at the time of the blood draw were typically over two years for non-twin sibling pairs (range = 0.08 - 5.38, *M* = 2.86, median = 2.59, *SD* = 1.56) and near zero for the MZ and DZ twin pairs (range = 0 - 0.52, *M* = 0.04, median = 0.00, *SD* = 0.10).

### DNA extraction

DNA was isolated from saliva, buffy coat, or peripheral blood mononuclear cells (PBMC) samples provided by the participants. Every tissue type was collected from every participant.

#### Saliva DNA Extraction

Saliva (2 mL) was collected from each participant in a 15 mL conical tube and a cotton swab containing dried reagents was added, bringing the solution to 1% SDS, 10 mM EDTA, 50 mM Tris pH 8.0 final concentration. The saliva was then stored at room temperature until DNA was extracted. The storage time ranged from about 3 years to 5 years. DNA was prepared from saliva using a protocol modified from Haberstick & Smolen (Haberstick & Smolen, 2004). Briefly, 600 µL of saliva in lysis buffer was sequentially treated with RNase A and proteinase K followed by precipitation with two volumes of isopropyl alcohol and pelleted, washed with 1mL 70% ethanol, air dried and resuspended in 110 µL of TE Buffer pH 8.0 (Invitrogen).

#### Buffy Coat DNA Extraction

Blood samples were collected by venipuncture into Monoject EDTA (K3) blood collection tubes (Covidien). The samples were processed following manufacturer instructions. Buffy coats were stored at −80C after processing on the day they were collected. DNA was isolated from 500 µL of thawed buffy coat using Maxwell 16 Blood DNA Purification Kits (Promega) on the Maxwell 16 instrument (Promega) following the manufacturer’s protocol.

#### Peripheral Blood Mononuclear Cell DNA Extraction

Blood samples were collected by venipuncture into Vacutainer CPT blood collection tubes (BD). The samples were processed following manufacturer instructions. PBMCs washed with 1x phosphate buffered saline were lysed in 1 mL lysis buffer (1% SDS, 50 mM Tris pH 8.0, 10 mM EDTA) and stored at −80C after processing on the day they were collected. DNA was prepared from PBMCs using a protocol modified from Haberstick & Smolen (Haberstick & Smolen, 2004). Briefly, PBMCs in lysis buffer were sequentially treated with RNase A and proteinase K followed by precipitation with two volumes of isopropyl alcohol and pelleted, washed with 1mL 70% ethanol, air dried and resuspended in 110 µL of TE Buffer pH 8.0 (Invitrogen).

The concentration of purified DNA from all three sources was quantified utilizing the Quant-iT PicoGreen dsDNA Assay Kit (Thermo Fisher Scientific) and then adjusted to 30-50 ng/µL in 1x TE. DNA methylation was measured on the EPIC v1 array (Illumina) by the Johns Hopkins University Genetic Resources Core Facility. In addition to the 273 samples (91×3), we submitted 3 additional technical duplicate samples of each tissue type for a total of 282 samples. DNA methylation of these technical duplicates was highly correlated within each tissue type (correlation, *r*, of methylation betas all ≥ 0.97); see Supplemental Figure S1).

### Data processing

Raw methylation IDAT files were imported and preprocessed using *minfi* v3.23 (Aryee et al., 2014; Fortin, Triche, & Hansen, 2017). We filtered poor quality probes using a detection p-value (*minfi::detectionP*) threshold of 0.05 and excluded probes with missingness ≥1% across samples (retaining 820,604 probes). We excluded samples with ≥1% missingness across probes, excluding 11 samples among 9 unique individuals, all of saliva origin (Supplemental Figure S2), a more stringent missingness threshold than other saliva-based analyses (Middleton et al., 2022). Using the *mapToGenome* and *dropLociWithSnps* functions, we removed any probes that mapped to known polymorphic sites, retaining 792,991 probes. We next performed background correction (*minfi::preprocessNoob*) and then normalization (*wateRmelon::dasen*) (Pidsley et al., 2013; Y. Wang et al., 2021). We assessed the degree of normalization using *wateRmelon::qual*, finding a greater normalization effect was required for the saliva samples relative to the buffy coat or PBMC tissue samples (Supplemental Figure S3). Methylation beta and M values were extracted using *getBeta* and *getM*. We further assessed overall differences by randomly selecting 5,000 probes and performing principal components on the M values (Supplemental Figure S4), finding clear, overall separation of the tissues, with a notable separation of saliva. Quality control led to the removal of 11 total saliva samples among 9 unique individuals, while all BC and PBMC samples passed quality control. After QC, we recalculated correlations of probes across the methylome for each pair of technical duplicates for samples that passed, with all *r* > 0.99. The final analysis sample sizes were 91 for both PBMC and BC, and 82 for saliva.

Cell proportions were estimated with *epidish* for blood tissues and *BeadSorted.Saliva.EPIC (Middleton et al., 2022)* using the Bioconductor *ExperimentHub* package (Morgan & Shepherd, 2025) for saliva samples, using the blood and saliva tissue references from each package, respectively (Supplemental Figure S5). Beta values were adjusted for cell type proportions and by plate.

### Analyses

DNA methylation clocks and the respective age acceleration residuals were calculated using the *dnaMethyAge* package in R entering beta values and the phenotype information of chronological age (Y. Wang et al., 2024) across all individuals. Age acceleration values were calculated using the default linear method where we retained residuals of methylation clock values regressed on chronological age. Sibling relatedness was not factored into computing the methylation age or age acceleration but was later accounted for in regression models and twin analyses. Within the *dnaMethyAge* package, calculations include 15 clocks such as Hannum, epiTOC, PhenoAge, PCGrimAge, and DunedinPACE (see Table 1 and Supplemental Table S1). One core clock of the *dnaMethyAge* package, cAge, was not included in the analysis due to a non-conformable argument run error. While we present findings on all 15 clocks, we prioritized 8 well-known clocks that included adults in clock training and produced DNA methylation ages or rate of aging as outcomes (see Table 1).

**Table 1.** Probe Coverage Across *dnaMethyAge* Clocks.

|  | DNAm Clock | Clock Type | Tissue(s) Trained | Total Probes | Probes Present | Percent Present | Weight Fraction Missing |
| --- | --- | --- | --- | --- | --- | --- | --- |
| 1 | CorticalClock <sup>a</sup> | 1 | Brain cortex | 348 | 345 | 99.1% | 0.13% |
| 2 | DunedinPACE <sup>b</sup> | 2 | Blood | 173 | 173 | 100.0% | NA |
| 3 | Hannum <sup>c</sup> | 1 | Whole blood | 71 | 65 | 91.5% | 8.97% |
| 4 | Horvath <sup>d</sup> | 1 | 27 tissues | 353 | 335 | 94.9% | 3.87% |
| 5 | Horvath2 <sup>e</sup> | 1 | 8 tissues | 391 | 382 | 97.7% | 1.38% |
| 6 | PCGrimAge <sup>f</sup> | 2 | Blood | 78,465 | 78,168 | 99.6% | NA |
| 7 | PhenoAge <sup>g</sup> | 2 | Whole blood | 514 | 509 | 99% | 0.35% |
| 8 | ZhangQ <sup>h</sup> | 1 | Blood, saliva | 515 | 509 | 98.8% | 0.22% |

*Notes.* DNAm Clock = DNA methylation clock available from *dnaMethyAge*; Clock type: 1=First-generation (trained on chronological age), 2=Next-generation (trained on health-related variables), 3=Biological processes clocks. <sup>a</sup> – (Shireby et al., 2020); <sup>b</sup> – (Belsky et al., 2022); <sup>c</sup> – (Hannum et al., 2013); <sup>d</sup> – (Horvath, 2013); <sup>e</sup> – (Horvath et al., 2018); <sup>f</sup> – (Higgins-Chen et al., 2022; Lu et al., 2019); <sup>g</sup> – (Levine et al., 2018); <sup>h</sup> – (Zhang et al., 2019).

We estimated rank-based Spearman correlations (*r*_*s*_) of methylation age among the sample mediums, as well as their correlations with chronological age, for each of the DNA methylation clocks. For each clock, we completed 100 iterations of drawing one random family member from each family set and calculated the rank-based Spearman correlation among sample mediums or with chronological age, to assess correlations without confounding familial factors. We selected Spearman correlation coefficients as effect sizes which are generally more robust to outliers or non-normality (de Winter, Gosling, & Potter, 2016).

To test differences in DNA methylation clocks by sample medium that were measured across the same individuals and individuals in the same family (sharing genetic and environmental influences), we fitted a restricted maximum likelihood mixed-effects model using the *nlme* package’s *lme()* function in R (Pinheiro, 2021). For each of the 15 different DNA methylation clocks, methylation age was the outcome for each model, with sample medium and chronological age at the time of the blood draw (centered at the mean chronological age, 30.90 years) as fixed effects. The unique family ID was modeled as a random effect, estimating the among-family standard deviation while the residual standard deviation quantified the within-family (individual) variance and any measurement error. Estimating the family random effects accounted for relatedness among participants.

In sensitivity analyses, we fitted mixed-effects models to DNA methylation clocks, omitting chronological age. We also fitted models to age acceleration estimates from *dnaMethyAge* (Y. Wang et al., 2024), which represents the residuals of methylation clock values regressed on chronological age. Thus, for those clocks that estimate a DNA methylation age, positive values for age acceleration measures index faster aging and negative values index slower aging than expected from chronological age.

### Twin comparisons

We calculated Spearman correlations within MZ and DZ twin pairs and created scatterplot visualizations. MZ twins are genetically identical, whereas DZ twins share half of their segregating genes on average; both MZ and DZ pairs may also be similar due to shared environments (Knopik, Neiderhiser, DeFries, & Plomin, 2017). Any person-specific environmental influences (non-shared between twin pairs) will contribute to MZ and DZ twin pair dissimilarity, as well as any measurement error. Hence, the MZ correlation represents the lower boundary of measurement reliability (Hagenbeek, van Dongen, Pool, & Boomsma, 2022).

Methylation age acceleration estimates were used for MZ and DZ twin pair comparisons of prioritized clocks where positive values index faster aging and negative values index slower aging, analogous to the DunedinPACE clock. For other clocks that that estimated biological processes rather than a methylation age (or transformation thereof), we retained the original absolute measures that reflect biological processes such as rates of aging or mortality. We calculated Spearman twin correlations randomizing twin member assignment over 100 iterations, where each twin was randomly assigned as twin1 or twin2 to provide a sense of variability in twin correlations given the number of pairs included. From these samples, we report the mean correlations and *SD* estimates for each tissue and clock, the 95% range of correlation estimates, and the percent of p<0.05 by clock and zygosity (Supplemental Table S5). To make comparisons by zygosity and clock generation, we applied a samplewise-adjusted weighted meta-analytic procedure to the obtained mean Spearman correlations and p-values using the DerSimonian–Laird option (Cheung & Chan, 2014) which relied on the *metafor* package (version 4.8-0) in R (Viechtbauer, 2010). This procedure allowed us to account for dependency in effect sizes between DNA methylation clocks by zygosity. Across the 15 clocks and three tissues, we report meta-analytic estimates of Spearman correlations, denoted *r̅*_*s*,*SAdj.wt*_, standard errors, and confidence intervals to compare MZ and DZ pairs where each zygosity group produced 45 estimates. We refer to unweighted mean correlations as *r̅*_*s*_ alongside their corresponding *SD* estimates.

All R code that was utilized from data processing to data analysis can be found using the GitHub link: https://github.com/rjbru/Methylation.git.

## Results

Metrics of all DNA methylation clocks successfully run using the *dnaMethyAge* package, their probe coverage, and the percentage of missing CpG probe weight relative to all clock CpG probe weights (Weight Fraction Missing) are shown in Table 1 and Supplemental Table S1. ‘Probes present’ represents coverage across all sample types (i.e., BC, PBMC, and saliva). Probe coverage of the ‘prioritized’ clocks were universally high (>91%), though other less-utilized clocks had much lower probe coverage (Table S1). Across all calculated clocks, the missing probes were assigned the mean values from each respective reference dataset in calculating trained phenotype outcomes by the *dnaMethyAge* package.

Average spearman rank correlations of DNA methylation clock and chronological ages using 100 different iterations where one random member from each family was used for the correlation analyses are shown in Figure 1 and Supplemental Figure S6, with the correlation ranges shown on Supplemental Table S2. Spearman rank correlations showed consistent, strong correlations between BC and PBMC methylation ages (*r̅*_*s*_ = 0.71, *SD* = 0.11; range = 0.50 – 0.93), whereas saliva showed weaker correlations with BC (*r̅*_*s*_ = 0.24, *SD* = 0.24, range = 0.06 – 0.93) and PBMC across most clocks (*r̅*_*s*_ = 0.25, *SD* = 0.25; range = −0.03 – 0.90). The highest average cross tissue associations by clocks (three correlations each) were PCGrimAge (*r̅*_*s*_ = 0.92, range = 0.90 – 0.93) followed by ZhangQ (*r̅*_*s*_ = 0.74, range = 0.61 – 0.88) then Horvath2 (*r̅*_*s*_ = 0.65, range = 0.51 – 0.81), reflecting next-generation and multi-tissue first-generation clocks. Other clocks showing moderate average associations include both first- and next-generation clocks (*r̅*_*s*_ = 0.47 – 0.54), for the Cortical Clock, DunedinPace, Hannum, Horvath, and PhenoAge.

**Figure 1.**
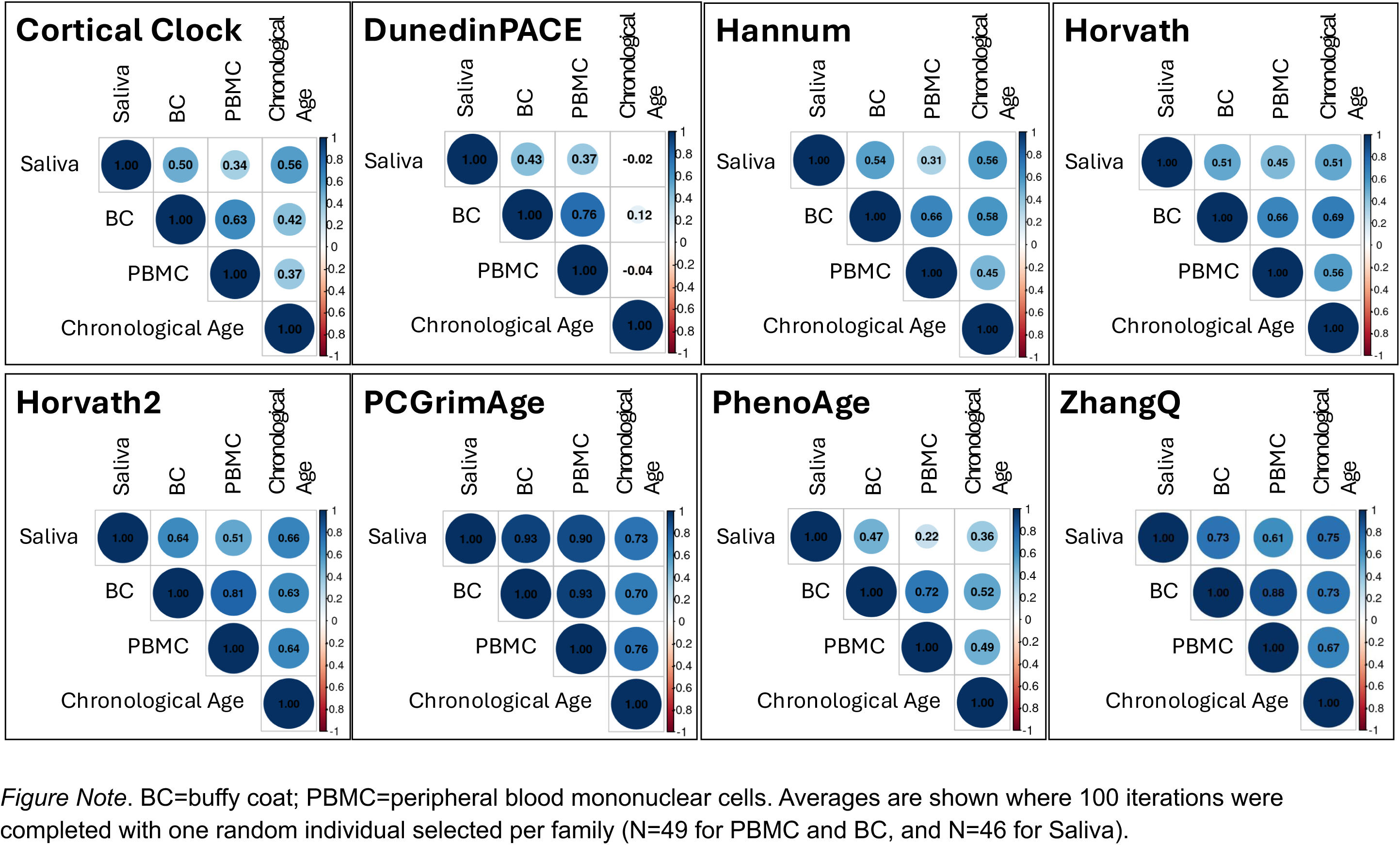
Spearman rank correlation averages among DNA methylation ages and with chronological age by sample medium.

In the random-effects model results using *lme*, which account for between-family effects, the estimated fixed effect intercept reflects the expected BC-based methylation age for clocks within the *dnaMethyAge* package. The *lme* results suggest that PBMC methylation age effect is not significantly different from the BC-based methylation age (i.e., the intercept). However, there was a significant effect of saliva tissue on DNA methylation clocks, with estimated older methylation clock values for 14 of 15 clocks (Table 2, Supplemental Tables S3-S4). For example, saliva-based ages were significantly different from the blood-based ages across blood-based DNA methylation clocks when considering Hannum (B_Saliva_ = 12.514, p = <0.001), Horvath (B_Saliva_ = 7.423, p = <0.001), Horvath2 (B_Saliva_ = 16.411, p = <0.001), PCGrimAge (B_Saliva_ = 9.102, p = <0.001), PhenoAge (B_Saliva_ = 19.723, p = <0.001) and ZhangQ (B_Saliva_ = 3.768, p = <0.001). Non-blood based DNA methylation clocks also showed significant differences for saliva including Cortical Clock (B_Saliva_ = 12.343, p = <0.001) and PedBE (B_Saliva_ = 10.680, p = <0.001). Multiple test correction using the Hommel method (Hommel, 1988) resulted in no difference in conclusions, where significant p-values remained across the 14 of 15 clocks (p = 1.253e-122 to 6.260e-20 for all but DunedinPACE which remained at p=0.486). Chronological age at the time of the blood draw was significantly positively associated with these aforementioned clocks, as expected, where one elapsed chronological year predicted 0.137 to 0.968 increased years in methylation age.

**Table 2.** Linear Mixed-effects Model Fit of Fixed Effects, DNA Methylation Clocks by Sample Medium.

| DNA Methylation Clock | B | SE | df | p-value | Between Family SD | Within Family SD |
| --- | --- | --- | --- | --- | --- | --- |
| 1. Cortical Clock |  |  |  |  |  |  |
| Intercept [BC=ref] | 38.023 | 0.369 | 212 | <0.001 | 1.662 | 2.682 |
| PBMC | -0.057 | 0.398 | 212 | 0.886 |  |  |
| Saliva | 12.343 | 0.410 | 212 | <0.001 |  |  |
| Chronological Age | 0.540 | 0.074 | 212 | <0.001 |  |  |
| 2. DunedinPACE |  |  |  |  |  |  |
| Intercept [BC=ref] | 1.304 | 0.011 | 212 | <0.001 | 0.057 | 0.070 |
| PBMC | 0.003 | 0.010 | 212 | 0.785 |  |  |
| Saliva | -0.007 | 0.011 | 212 | 0.486 |  |  |
| Chronological Age | 0.003 | 0.002 | 212 | 0.224 |  |  |
| 3. Hannum |  |  |  |  |  |  |
| Intercept [BC=ref] | 36.023 | 0.365 | 212 | <0.001 | 1.606 | 2.692 |
| PBMC | 0.136 | 0.399 | 212 | 0.734 |  |  |
| Saliva | 12.514 | 0.411 | 212 | <0.001 |  |  |
| Chronological Age | 0.645 | 0.072 | 212 | <0.001 |  |  |
| 4. Horvath |  |  |  |  |  |  |
| Intercept [BC=ref] | 21.830 | 0.471 | 212 | <0.001 | 2.117 | 3.417 |
| PBMC | 0.386 | 0.507 | 212 | 0.447 |  |  |
| Saliva | 7.423 | 0.522 | 212 | <0.001 |  |  |
| Chronological Age | 0.797 | 0.094 | 212 | <0.001 |  |  |
| 5. Horvath2 |  |  |  |  |  |  |
| Intercept [BC=ref] | 18.420 | 0.396 | 212 | <0.001 | 1.699 | 2.963 |
| PBMC | 0.085 | 0.439 | 212 | 0.847 |  |  |
| Saliva | 16.411 | 0.452 | 212 | <0.001 |  |  |
| Chronological Age | 0.811 | 0.078 | 212 | <0.001 |  |  |
| 6. PCGrimAge |  |  |  |  |  |  |
| Intercept [BC=ref] | 48.121 | 0.289 | 212 | <0.001 | 1.824 | 1.169 |
| PBMC | 0.135 | 0.173 | 212 | 0.436 |  |  |
| Saliva | 9.102 | 0.179 | 212 | <0.001 |  |  |
| Chronological Age | 0.925 | 0.058 | 212 | <0.001 |  |  |

| 7. PhenoAge |  |  |  |  |  |  |
| --- | --- | --- | --- | --- | --- | --- |
| Intercept [BC=ref] | 20.216 | 0.701 | 212 | <0.001 | 2.760 | 5.594 |
| PBMC | 0.269 | 0.829 | 212 | 0.746 |  |  |
| Saliva | 19.723 | 0.854 | 212 | <0.001 |  |  |
| Chronological Age | 0.899 | 0.136 | 212 | <0.001 |  |  |
| 8. ZhangQ |  |  |  |  |  |  |
| Intercept [BC=ref] | 26.164 | 0.346 | 212 | <0.001 | 1.643 | 2.411 |
| PBMC | 0.217 | 0.357 | 212 | 0.545 |  |  |
| Saliva | 3.768 | 0.368 | 212 | <0.001 |  |  |
| Chronological Age | 0.968 | 0.070 | 212 | <0.001 |  |  |
| Notes. BC = buffy coat; PBMC = peripheral blood mononuclear cells; SD = standard deviation. |  |  |  |  |  |  |

The DunedinPACE clock measure is scaled as a rate of aging (Belsky et al., 2022) with expected values of 1.0 if individuals’ rates are on par with chronological year; values >1.0 suggesting accelerating rates of aging and values less than 1.0 index a slower rate of aging. The average intercept suggested accelerated aging across the sample (B_intercept_ = 1.304, p = <0.001). However, there were no significant differences in aging rates of PBMC- versus BC-based estimates (B_PBMC_= 0.003, p = 0.785) or saliva-based versus BC-based estimates (B_Saliva_= −0.007, p = 0.486), supporting overall comparability. Chronological age was likewise non-significant (B_ChronologicalAge_ = 0.003, p = 0.224).

Visualizations of the DNA methylation clock values are shown for each of the three sample types (see Supplemental Figure S7), selecting one member of each sibling or twin pair, and all unpaired individuals (N=49 PBMC and BC, N=46 saliva). As demonstrated in the *lme* analyses, PBMC- and BC-based estimates have comparable mean DNA methylation ages and show similar distributions for the prioritized clocks, whereas saliva DNA methylation ages are older. Likewise demonstrated in the *lme* analyses, a number of first- and next-generation clocks show higher estimated methylation ages across tissues, e.g., Cortical Clock, Hannum, PCGrimAge. Both Horvath clocks and PhenoAge had younger methylation ages compared to chronological age, especially BC and PBMC. ZhangQ showed the most comparable age distribution to our sample and across three tissues.

Sensitivity analyses of all the 15 DNA methylation clocks omitted chronological age and achieved comparable results (see Supplemental Table S4), again revealing a distinction between saliva versus PBMC and buffy coat samples apart from the DunedinPACE clock. Models fitted to age acceleration residuals likewise show the distinction between saliva versus PBMC and buffy coat samples in 14 of the 15 clocks (see Supplemental Table S4), apart from the DunedinPACE as expected.

### Twin pair comparisons

We present Spearman rank correlations for all 15 DNA methylation clocks in Supplemental Table S5 by zygosity. Correlations were calculated using age-acceleration estimates for all first-generation clocks and next-generation clocks producing DNA methylation age estimators. For next-generation clocks reporting aging rates or mortality risk, and clocks tracking other biological processes, we report correlations using unadjusted DNA methylation clock estimates. To summarize the findings, we report meta-analytic estimates in Supplemental Table S5 and provide example visualizations in Figure 2.

**Figure 2.**
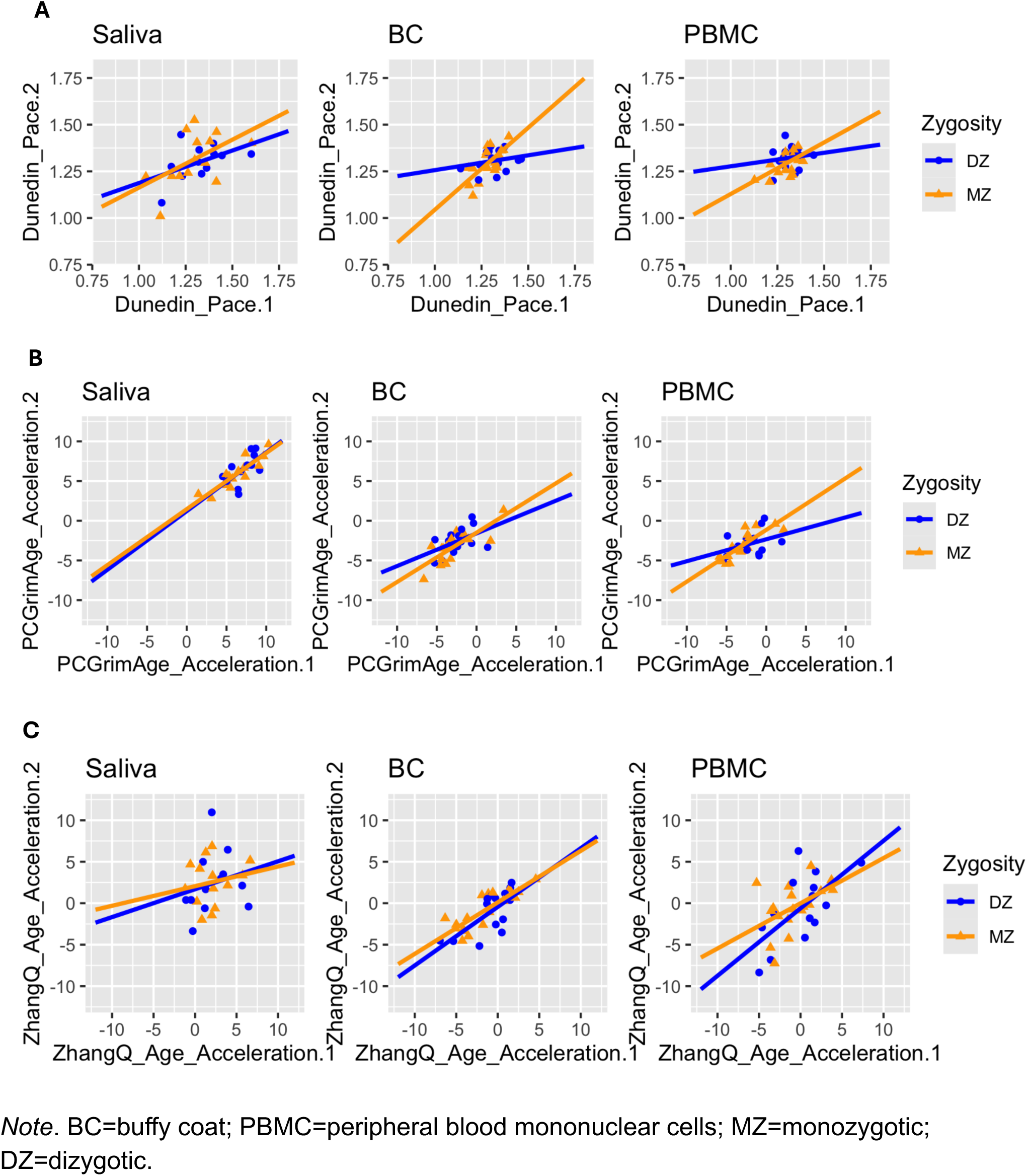
Scatterplots of Twin pair Age Acceleration estimates, separated by zygosity and sample medium: (A) Dunedin Pace, (B) PCGrimAge clock acceleration, (C) ZhangQ clock acceleration.

Across all clocks and tissue types tested here, MZ pairs were more correlated (*r̅*_*s*,*SAdj.wt*_=0.49, *se*=0.09, *p*<0.001, 95%CI=0.32,0.66) than DZ pairs (*r̅*_*s*,*SAdj.wt*_=0.38, *se*=0.07, *p*<0.001, 95%CI=0.25,0.52), but with overlapping confidence intervals. However, MZ pairs showed significant heterogeneity in correlations across tissue type (Cochran’s *Q*=6.77, *p*=0.034; *I*^2^=70.45%) with lower and more inconsistent correlations for saliva (*r̅*_*s*_=0.33, *SD*=0.25) and stronger more consistent correlations for BC (*r̅*_*s*_=0.64, *SD*=0.10) and PBMC (*r̅*_*s*_=0.49, *SD*=0.18). DZ pairs did not show significant heterogeneity (Cochran’s *Q*=0.70, *p*=0.703; *I*^2^=0). This suggests greater reliability of blood-based clocks as the strength of the MZ correlation represents a lower bound of reliability (Hagenbeek et al., 2022).

Meta-analysis of twin pair correlations by clock type suggested that the next-generation age acceleration estimates (PCGrimAge, PhenoAge, DunedinPACE, ZhangY) report moderately higher MZ correlations (*r̅*_*s*,*SAdj.wt*_=0.58, *se*=0.05, *p*<0.001, 95%CI=0.49,0.68) than DZ correlations (*r̅*_*s*,*SAdj.wt*_=0.40, *se*=0.09, *p*<0.001, 95%CI=0.23,0.58), but with overlapping confidence intervals. Notably, among all 15 clocks, the next-generation PCGrimAge clock showed the highest average MZ correlation (*r̅*_*s*_=0.85, *SD*=0.05) (see Supplemental Table S5). PCGrimAge showed the fifth highest average DZ correlation across all three tissue types (*r̅*_*s*_=0.48, *SD*=0.12). ZhangQ clock was stronger among DZ pairs but with greater variability than PCGrimAge (*r̅*_*s*_=0.56, *SD*=0.16). PCGrimAge and DunedinPace showed the largest MZ versus DZ differences in average correlations across all three tissues.

First-generation clocks show similar moderate correlations among MZ (*r̅*_*s*,*SAdj.wt*_=0.45, *se*=0.13, *p*<0.001, 95%CI=0.19,0.70) and DZ twins (*r̅*_*s*,*SAdj.wt*_=0.42, *se*=0.06, *p*<0.001, 95%CI=0.31,0.52). Similar correlation patterns were observed for recent multi-tissue clocks (all first-generation, excluding Horvath, 2013), where MZ pairs (*r̅*_*s*,*SAdj.wt*_=0.49, *se*=0.15, *p*<0.002, 95%CI=0.19,0.79) and DZ pairs (*r̅*_*s*,*SAdj.wt*_=0.45, *se*=0.09, *p*<0.001, 95%CI=0.26,0.63) were moderately correlated with overlapping confidence intervals.

In Figure 2, we illustrate a scatterplot of twin pair similarity for age acceleration metrics for 26 to 32 twin pairs (14 - 18 MZ pairs, 12 - 14 DZ pairs), displaying DunedinPACE, PCGrimAge and ZhangQ age acceleration estimates, where twin 2’s age acceleration is plotted against twin 1’s age acceleration (with twin assignment random). For added visual comparison, we plot inferred regression lines where the steepness of the line indexes the relative similarity of MZ versus DZ twins. For DunedinPACE, the MZ regression line is visually steeper than the DZ pairs suggesting greater similarity for all three tissues. For PCGrimAge, differences are more evident for BC and PBMC than saliva (and saliva notably shows positive age accelerations relative to PBMC and BC). For the ZhangQ age acceleration, the inferred regression lines for MZ pairs are comparable to DZ pairs suggesting little difference by zygosity, for all three tissues.

## Discussion

We evaluated methylation ages across different tissues to gauge the equivalence of blood and saliva tissues commonly sampled and biobanked. Strong correlations of methylation ages with chronological age were observed across all tissue types. However, blood-based PBMC and BC methylation age estimates were more correlated with each other than with saliva-based estimates. Saliva-based methylation ages were significantly older relative to the blood-based DNA methylation ages, whereas no significant differences were found between PBMC and BC methylation ages. The next-generation age-acceleration clock, DunedinPACE, was the exception showing no tissue type differences. Overall, while blood-based methylation ages may be younger, correlational analyses with chronological age suggest that saliva doesn’t perform any worse. Patterns of MZ similarity were stronger for blood-based clocks than saliva-based clocks, while DZ pairs showed consistent similarity across tissue type. MZ pairs showed modestly greater similarity than DZ pairs, particularly for next-generation clocks. Three clocks showed notable cross-tissue consistency or comparability across analyses: the first-generation multi-tissue ZhangQ, and the next-generation DunedinPace and PCGrimAge clocks, generally supporting our expectations, albeit with some caveats.

Our results are among the first to demonstrate that DNA methylation clock ages do not significantly differ between BC and PBMC in a larger sample of adults. Moreover, BC and PBMC based clocks more closely aligned with each other than with saliva based clocks. Prior work (Apsley et al., 2025) evaluated BC in children only and PBMC in adults only but generally observed higher methylation ages in oral-tissues than blood-based tissues, similar to our work. While the Skin and Blood (Horvath2) clock exhibited the greatest concordance in terms of tissue comparability in this prior work (Apsley et al., 2025), the authors noted older methylation ages for saliva versus PBMC. Our results for PhenoAge and PCGrimAge DNA methylation ages and accelerations are consistent with Apsley et al. (2025), again with older methylation ages and accelerations for saliva. The lower cross-tissue correlation of saliva-based clocks with BC and PBMC is consistent with a small study of 16 individuals focused on EPIC v1 versus EPIC v2 platform comparability (c.f., supplemental data EPIC v1,Tay et al., 2025) and between PBMC and BC with buccal cells and saliva (Apsley et al., 2025). Notably, however, in our current study the next-generation DunedinPACE age acceleration clock showed no significant mean differences between biological mediums, generally consistent with Zarandooz and Raffington (2025) but unlike (Apsley et al., 2025) which showed higher accelerations in oral-based versus blood-based tissues. Indeed, Zarandooz and Raffington (2025; c.f., Supplemental Figure 1) show little mean differences across any of the age acceleration estimates across clocks although their primary statistical analysis evaluated intraclass correlations (ICCs). Moreover, they observed moderate cross-tissue association between whole blood and saliva for DunedinPACE (ICC=0.68, CI= 0.57–0.79) compared to our associations (*r̅*_*s*_ = 0.37 – 0.43, for saliva-PBMC and saliva-BC, respectively).

We expected that first-generation multi-tissue clocks and next-generation clocks would show greater associations across sample mediums, with the Horvath2, ZhangQ, and PCGrimAge, clocks showing strong cross-tissue correlations compared to Cortical, Hannum, and Horvath first-generation clocks which showed moderate correlations. Our results are consistent in principle with Apsley et al’s (2025) findings for the Skin and Blood (Horvath2) clock and Zarandooz and Raffington (2025) findings for PCGrimAge although neither Apsley et al (2025) nor Zarandooz and Raffington (2025) tested ZhangQ. Moreover, for each biological tissue that we tested, the multi-tissue ZhangQ and next-generation PCGrimAge methylation ages showed the strongest correlations with chronological age. Similarly, Zarandooz and Raffington (2025) reported stronger correlations of whole blood-versus saliva-based methylation age accelerations comparing next-generation clocks to first-generation clocks in a recent study of 107 individuals aged 5 to 74 years. Indeed, they observed that PCGrimAge showed the strongest intraclass correlations for blood and saliva. However, we observed that absolute DNA methylation ages and age accelerations were divergent across clocks, except for DunedinPace. Hannum, trained on whole blood (Hannum et al., 2013), and Cortical Clock showed notably higher estimated methylation ages compared to chronological age compared to other clocks for blood-based BC and PBMC. Both Horvath clocks and PhenoAge had younger methylation ages compared to chronological age across all samples, but noticeably so in the BC and PBMC. The ZhangQ clock showed more consistent ages across all three tissues and was closest to our sample’s distribution of chronological ages. Age-acceleration residuals were not a panacea given that saliva routinely showed higher age-accelerations than other tissues, apart from DunedinPace as noted.

Twin similarity patterns were commensurate with the overall findings, showing patterns of greater pair similarity for blood-based than saliva-based clocks. MZ pairs were on average more similar than DZ pairs for second-generation than first-generation clocks across tissue type, with patterns largely consistent with publications on clock-related CpGs and second-generation clocks (Jylhava et al., 2019; Reynolds et al., 2020). DZ pair correlations appear numerically higher than MZ pair correlations in a few instances (e.g., CorticalClock Saliva, Hannum PBMC, Horvath PBMC); however, these differences were not formally tested and additional considerations such as sampling variability, testing clocks trained on other tissue types, and a small sample size are noteworthy. In the current work, the greatest divergence in MZ and DZ similarity was observed for the next-generation PCGrimAge age acceleration (Higgins-Chen et al., 2022; Lu et al., 2019) and DunedinPACE (Belsky et al., 2022), consistent with genetic influences on the pace of biological aging. Nonetheless, larger sample sizes are needed to estimate formal biometrical estimates of heritability and environmentality and is part of our future work in CATSLife, where this current study will guide our tissue sample selection among 1160 participants in CATSLife1 and CATSLife2 (ongoing).

While there is no perfectly performing clock in our analysis, three showed notable consistency or comparability. First, the multi-tissue ZhangQ clock showed the most comparable DNA methylation ages across tissues compared to the chronological ages in our sample, albeit saliva was significantly older/accelerated, plus ZhangQ evidenced generally moderate MZ and DZ twin correlations. The next-generation DunedinPace clock showed comparable aging rates across three tissues and moderate to strong MZ similarity with divergent MZ and DZ twin differences, as expected under biometrical theory for additive genetic influences (Knopik et al., 2017). The PCGrimAge clock, though showing older DNA methylation ages, demonstrated strong cross-tissue correlations, and strong MZ and DZ similarity with divergent MZ and DZ twin differences. Overall, further insights into the methylation age and age-acceleration differences, for both blood-based and saliva mediums are needed. Age-related CpG sites specific to saliva are of increasing interest (Xiao et al., 2025), and calls for cell-specific work are advancing which may produce clocks that work across tissue types that share common cell types (Teschendorff & Horvath, 2025).

In choosing between tissue types for methylation analysis, it is important to evaluate the proximity of blood versus saliva tissues to biological pathways of interest. While it might be presumed that blood-based methylation is more proximal than saliva-based methylation to brain-related mechanisms, saliva may be a reasonable medium given it likewise contains immune cell types (Theda et al., 2018). Moreover, biomarkers related to neurodegenerative outcomes, including p-tau and beta-amyloid components, are detectable in salivary tissues (Nijakowski, Owecki, Jankowski, & Surdacka, 2024). A caution is that while saliva is an attractive medium, in our study saliva samples were generally of a lower quality with nine samples removed during the QC steps. The differences in sample storage (−80C for buffy coats and PBMC lysate and room temperature for saliva for up to five years) could explain this difference in quality. As previous work has discussed (Staunstrup et al., 2017), overall DNA quality and potential contamination can be factors in utilizing saliva samples for methylation analysis. Some biobanks and longitudinal studies have saliva samples stored at room temperature (Spinola et al., 2025), and while we note cautions, collectively our work and others (Apsley et al., 2025; Zarandooz & Raffington, 2025) show promise of moderate to strong cross-tissue associations with select multi-tissue and next-generation clocks warranting further tool development.

It is important to note that our study is not without limitations. Our sample set includes individuals within established adulthood and where chronological age ranges were less than 15 years apart. Saliva had generally stronger effects of normalization, suggesting sample quality may be poorer in our saliva samples, although probe missingness for clock CpG sites (for prioritized clocks) was universally low for samples that passed QC thresholds. All retained samples passed our stringent missingness thresholds (stronger than other saliva-based analyses (Middleton et al., 2022); however, we cannot rule out that the differences in clocks estimated from saliva samples result only from or are affected by sample quality. Samples from PBMC and BC, however, did not suffer from this same issue, and were not statistically different in DNA methylation clock estimates (Table 2, Supplemental Table S3). We could not run cAge in our analysis due to a non-conformity error likely resulting from some CpG site missingness. Three additional clocks (Pan-Mammalian2, Pan-Mammalian3, and ZhangY, none among the prioritized clocks) had a large proportion of missing probes representing notable effect sizes for these outputs. While the *dnaMethyAge* package does inpute the mean value of the reference dataset across missing probes, the results of these three clocks should be interpreted with caution. Also, while we show notable differences between saliva compared to PBMC and BC, additional blood-based and other biological mediums may be more appropriate for methylation analysis. Future work including additional biological mediums, wider chronological age ranges of participants, longitudinal comparisons, and addressing cell-specific patterns, would each add to a greater understating of methylation profiles.

Our study offers insights on the comparability of biological tissues. While saliva is a less invasive method of collecting a biological medium for methylation, the costs of potential losses of samples due to poorer quality should be factored into decisions. Additional considerations include the consistently elevated saliva-based methylation ages predicted across individuals compared to blood-based methylation results. Three clocks showed notable, albeit imperfect, consistency or comparability across all three tissues: the first-generation multi-tissue ZhangQ, and the next-generation DunedinPace and PCGrimAge clocks, warranting further DNA methylation clock development. Broadly, our results support comparability of BC-based methylation to PBMC-based methylation and thus for banked buffy coat samples it may be an effective medium to address blood-based methylation when PBMC tissues are not available.

## Data availability

Individual level methylation data and all data used in this study have been submitted to NIAGADS (NG00145.v1; https://dss.niagads.org/datasets/ng00145/), where participant directives have allowed.

