## Supplemental Figures for "Differential DNA methylation clock ages across buffy coat (BC), peripheral blood mononuclear cells (PBMC), and saliva in individuals in early-to-mid adulthood"

**Figure S1.** Comparisons of methylation betas among technical duplicates, prior to QC.

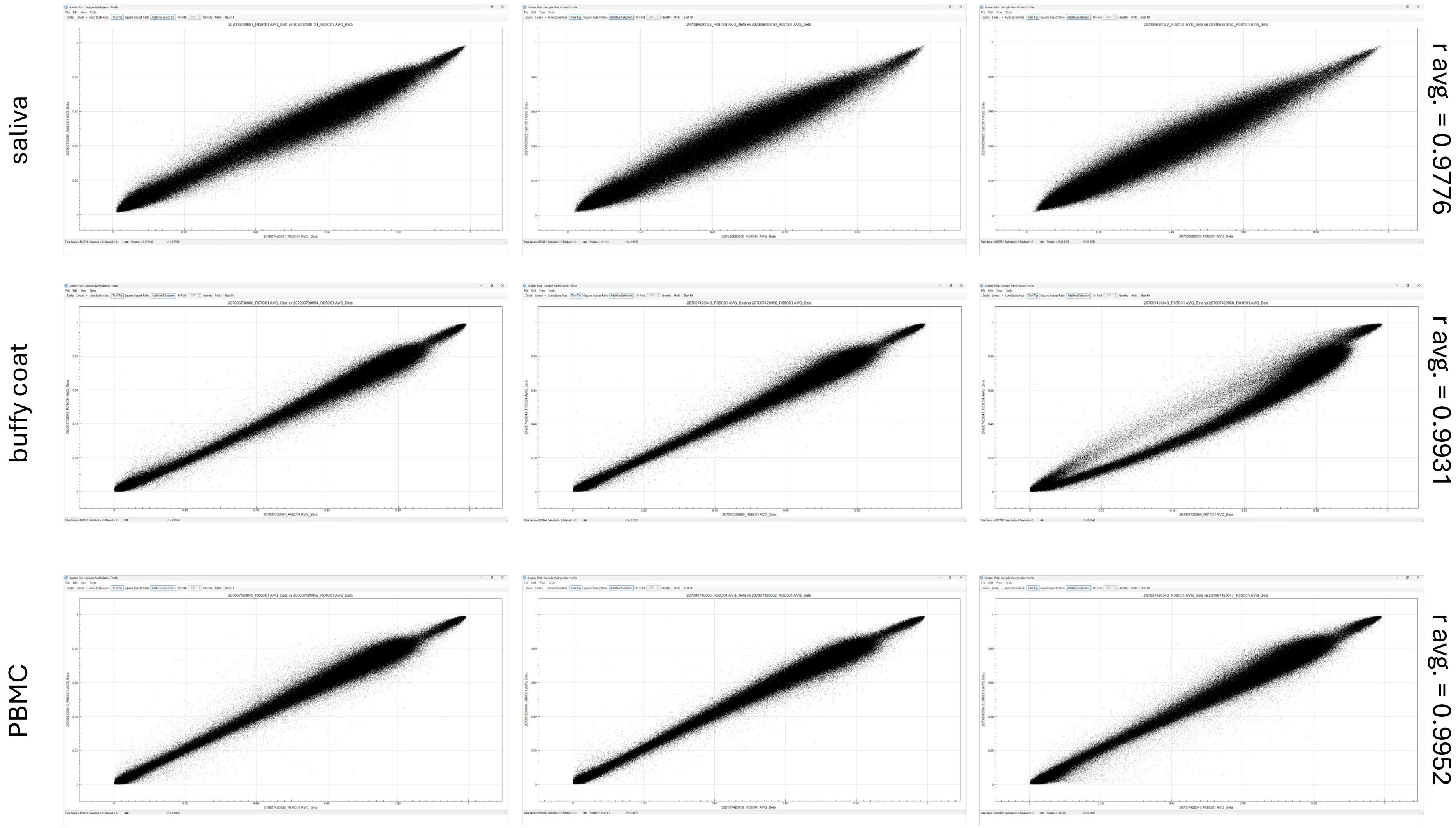

**Figure S2.** Distributions of sample (top left) and probe missingness (top right), including the numbers of individuals and probes excluded and retained by each. The 11 samples were all of saliva origin. Note that two of the excluded samples were duplicates, so the total number of unique individuals excluded was 9. Bottom: Sample missingness split by tissue.

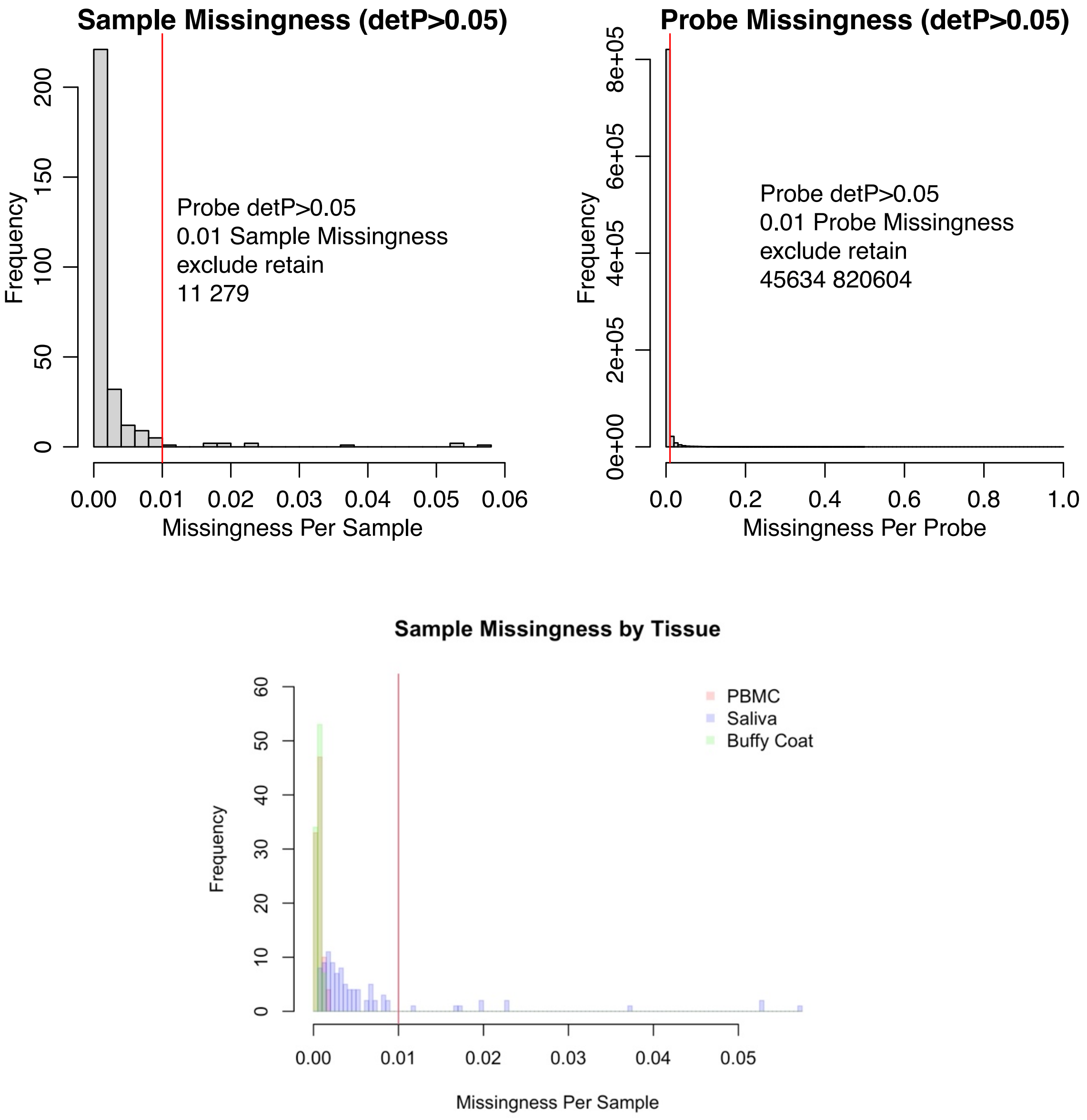

**Figure S3.** Normalization metrics using *wateRmelon*::*qual* by tissue. The normalization effect was strongest in the retained (Supplemental Fig. 1) Saliva samples, but similar between Buffy Coat and PBMC tissues samples. Different measures of normalization effect were similar.

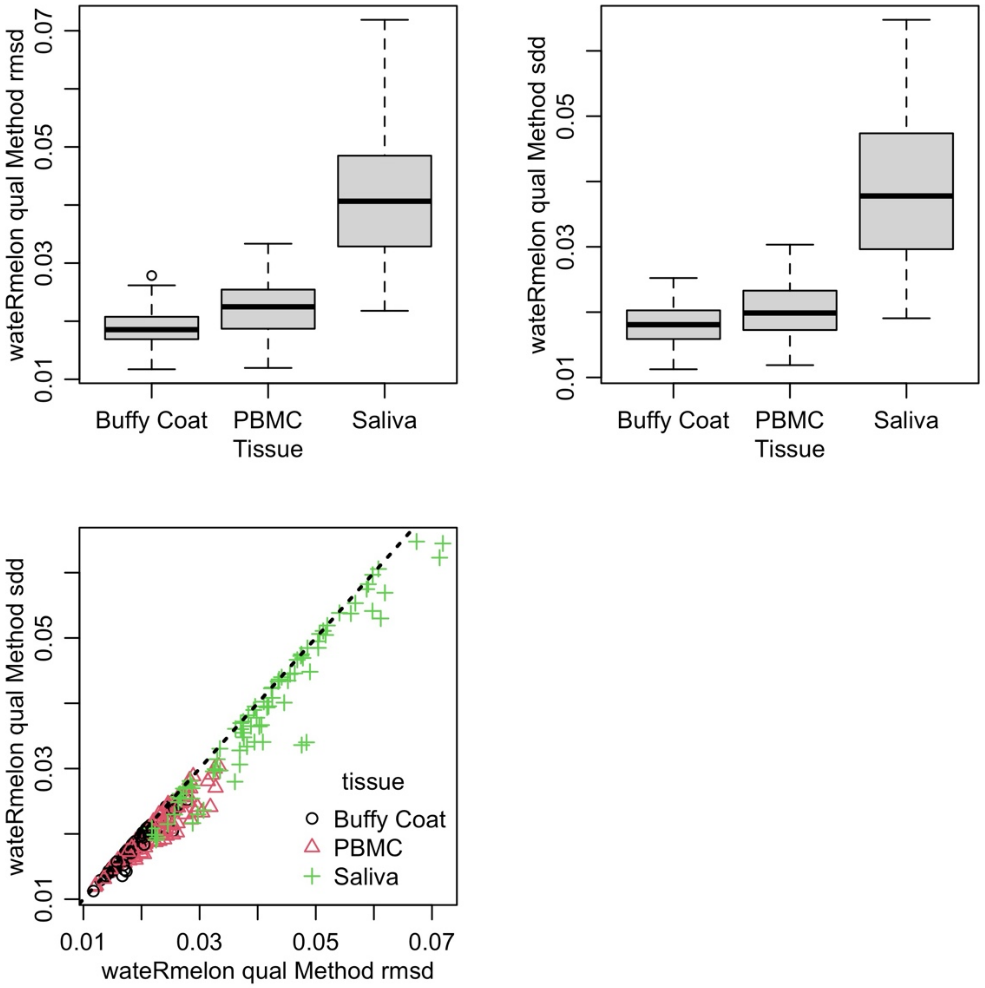

**Figure S4.** Principal component analysis of M values of 5,000 randomly selected probes across tissues.

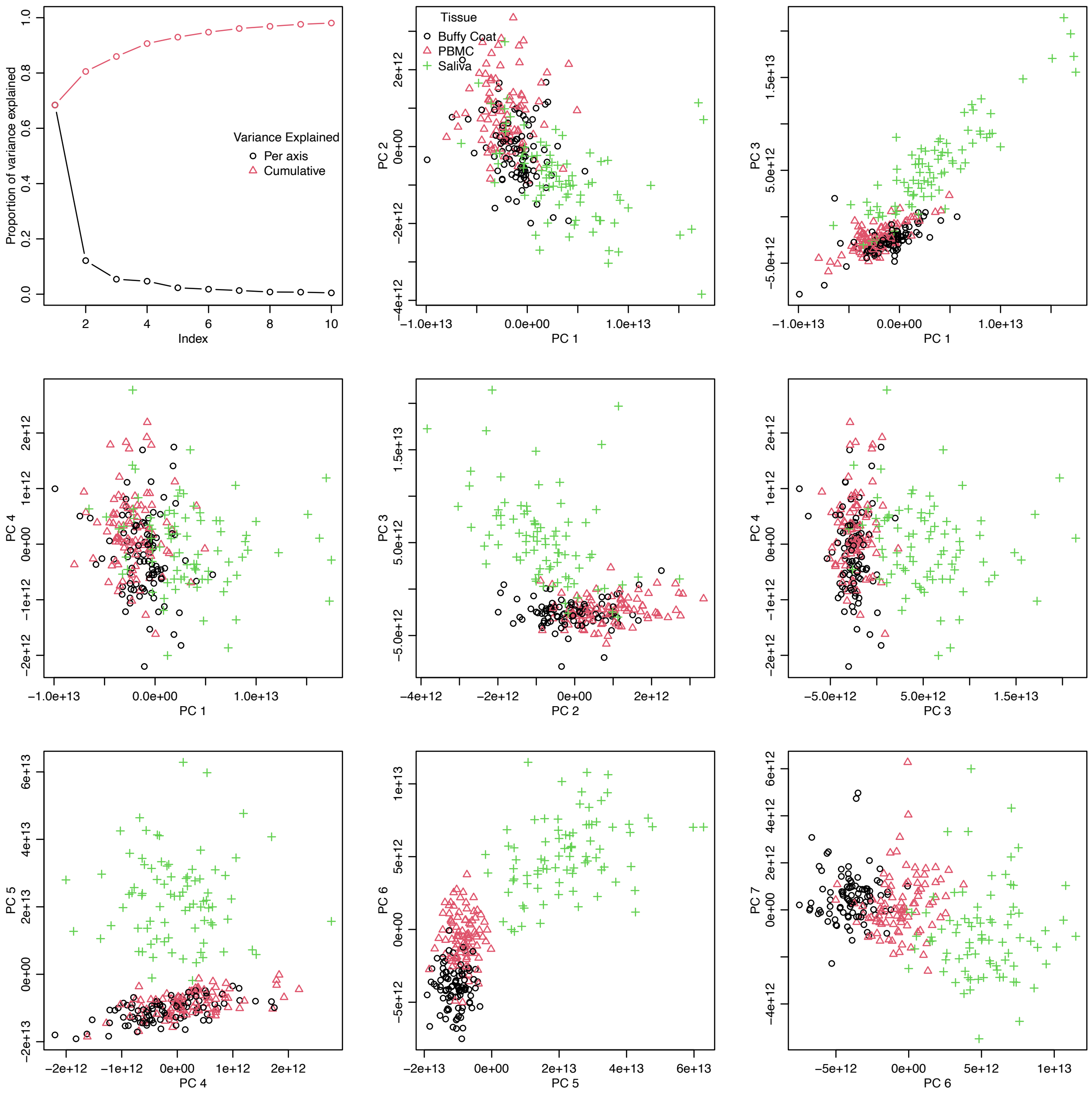

**Figure S5.** Cell type proportions estimated in the three tissue types, using the R packages *epidish* for blood and *BeadSorted.Saliva.EPIC* for saliva.

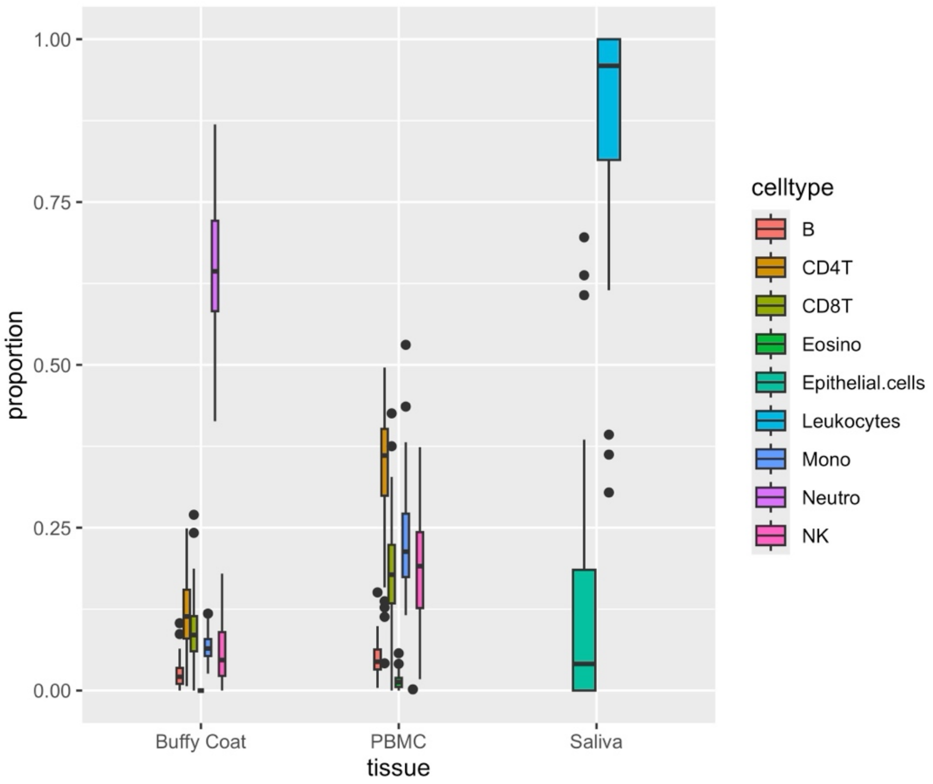

**Figure S6.** Spearman rank correlation averages among supplemental DNA methylation clocks with chronological age by sample medium.

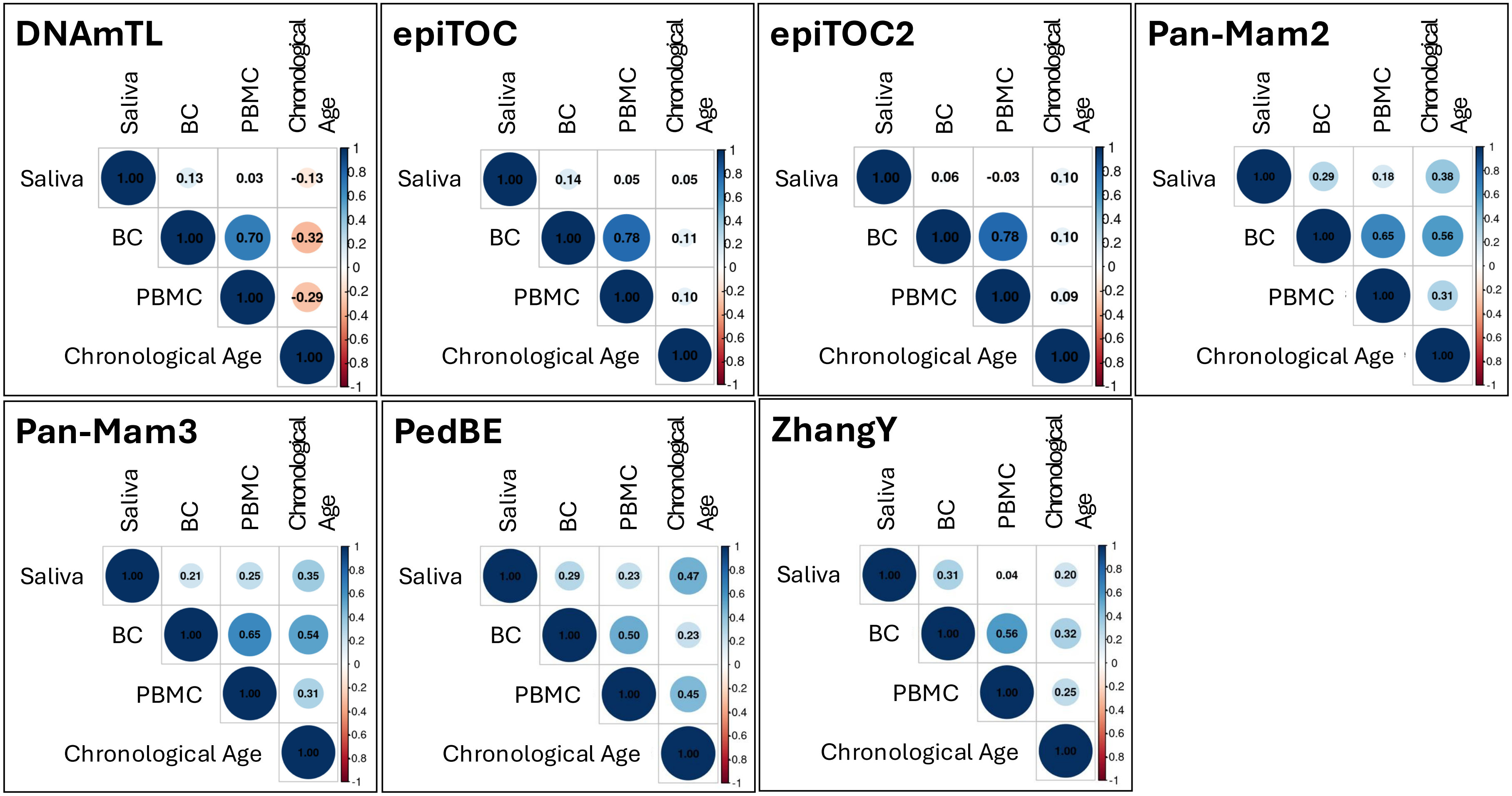


*Figure Note*. BC = buffy coat; PBMC = peripheral blood mononuclear cells

**Figure S7.** DNA methylation clock estimates by sample medium: (A) prioritized clocks, (B) Additional DNA methylation clocks.

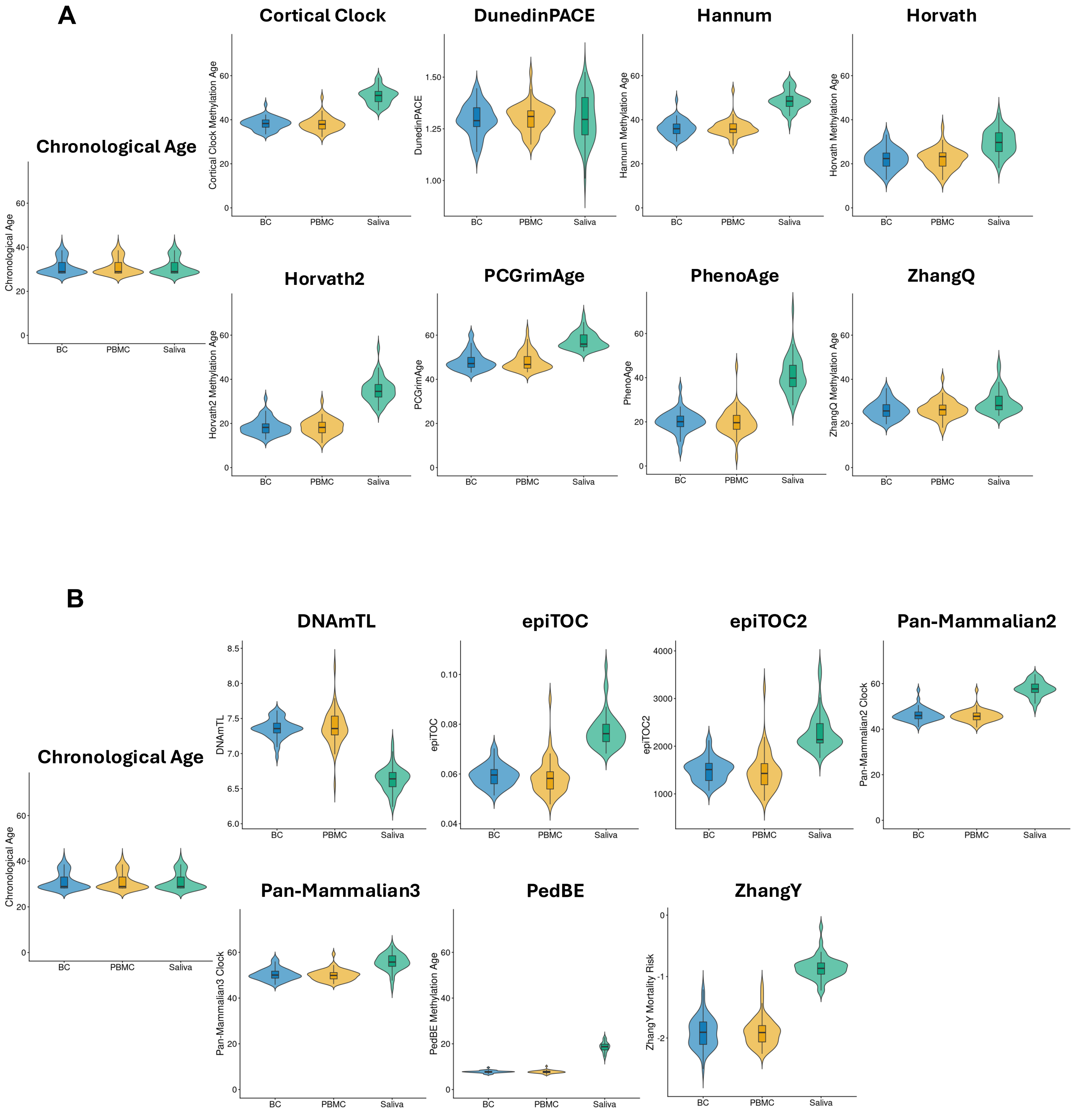


*Figure Note*. BC=buffy coat; PBMC=peripheral blood mononuclear cells. One random individual selected per family (N=49 for PBMC and BC, and N=46 for Saliva).
