## Supplemental Tables for "Differential DNA methylation clock ages across buffy coat (BC), peripheral blood mononuclear cells (PBMC), and saliva in individuals in early-to-mid adulthood"

**Table S1** - Probe Coverage Across *dnaMethyAge* Clocks.

| | DNAm Clock | Clock Type | Tissue(s) Trained | Total Probes | Probes Present | Percent Present | Weight Fraction Missing |
|---|---|---|---|---|---|---|---|
| 9 | DNAmTL [i] | 3 | Whole blood | 141 | 137 | 97.2% | 1.48% |
| 10 | epiTOC [j] | 3 | Whole blood | 385 | 352 | 91.4% | 7.90% |
| 11 | epiTOC2 [k] | 3 | Whole blood | 163 | 149 | 91.4% | 8.60% |
| 12 | Pan-Mammalian Clock2 [l] | 1 | 59 tissues,185 mammalian species | 816 | 167 | 20.5% | 73.99% |
| 13 | Pan-Mammalian Clock3 [l] | 1 | 59 tissues,185 mammalian species | 760 | 152 | 20.0% | 75.43% |
| 14 | PedBE [m] | 1 | Buccal epithelial cells | 95 | 93 | 97.9% | 2.26% |
| 15 | ZhangY [n] | 2 | Whole blood | 10 | 8 | 80% | NA |

*Notes*. DNAm Clock = DNA methylation clock available from *dnaMethyAge;* Clock type: 1=First-generation (trained on chronological age), 2=Next-generation (trained on health-related variables), 3=Biological processes clocks.
[i] – (Lu et al., 2019); [j] – (Yang et al., 2016); [k] – (Teschendorff, 2020); [l] – (Lu et al., 2023); [m] – (McEwen et al., 2020); [n] – (Zhang et al., 2017)

**Table S2** - Spearman rank correlation ranges (100 iterations drawing one random family member from each family) among DNA methylation ages and with chronological age by sample medium.

| CorticalClock | Saliva | BC | PBMC | Pheno Age | |
|---|---|---|---|---|---|
| Saliva | 1.00 | 0.70 | 0.58 | 0.65 | Maxes |
| BC | 0.28 | 1.00 | 0.81 | 0.56 | |
| PBMC | 0.07 | 0.31 | 1.00 | 0.57 | |
| Pheno Age | 0.43 | 0.27 | 0.17 | 1.00 | |
| | Mins | | | | |

| DunedinPACE | Saliva | BC | PBMC | Pheno Age | |
|---|---|---|---|---|---|
| Saliva | 1.00 | 0.58 | 0.53 | 0.16 | Maxes |
| BC | 0.23 | 1.00 | 0.87 | 0.32 | |
| PBMC | 0.15 | 0.57 | 1.00 | 0.15 | |
| Pheno Age | -0.17 | -0.09 | -0.24 | 1.00 | |
| | Mins | | | | |

| Hannum | Saliva | BC | PBMC | Pheno Age | |
|---|---|---|---|---|---|
| Saliva | 1.00 | 0.68 | 0.52 | 0.67 | Maxes |
| BC | 0.37 | 1.00 | 0.80 | 0.68 | |
| PBMC | 0.05 | 0.45 | 1.00 | 0.60 | |
| Pheno Age | 0.41 | 0.46 | 0.27 | 1.00 | |
| | Mins | | | | |

| Horvath | Saliva | BC | PBMC | Pheno Age | |
|---|---|---|---|---|---|
| Saliva | 1.00 | 0.65 | 0.61 | 0.66 | Maxes |
| BC | 0.35 | 1.00 | 0.77 | 0.78 | |
| PBMC | 0.27 | 0.49 | 1.00 | 0.70 | |
| Pheno Age | 0.38 | 0.56 | 0.36 | 1.00 | |
| | Mins | | | | |

| Horvath2 | Saliva | BC | PBMC | Pheno Age | |
|---|---|---|---|---|---|
| Saliva | 1.00 | 0.78 | 0.72 | 0.78 | Maxes |
| BC | 0.50 | 1.00 | 0.89 | 0.72 | |
| PBMC | 0.29 | 0.70 | 1.00 | 0.77 | |
| Pheno Age | 0.48 | 0.54 | 0.36 | 1.00 | |
| | Mins | | | | |

| PCGrimAge | Saliva | BC | PBMC | Pheno Age | |
|---|---|---|---|---|---|
| Saliva | 1.00 | 0.95 | 0.93 | 0.78 | Maxes |
| BC | 0.89 | 1.00 | 0.96 | 0.81 | |
| PBMC | 0.83 | 0.88 | 1.00 | 0.82 | |
| Pheno Age | 0.68 | 0.59 | 0.66 | 1.00 | |
| | Mins | | | | |

| PhenoAge | Saliva | BC | PBMC | Pheno Age | |
|---|---|---|---|---|---|
| Saliva | 1.00 | 0.65 | 0.49 | 0.59 | Maxes |
| BC | 0.22 | 1.00 | 0.85 | 0.72 | |
| PBMC | -0.07 | 0.59 | 1.00 | 0.72 | |
| Pheno Age | 0.20 | 0.33 | 0.26 | 1.00 | |
| | Mins | | | | |

| ZhangQ | Saliva | BC | PBMC | Pheno Age | |
|---|---|---|---|---|---|
| Saliva | 1.00 | 0.85 | 0.75 | 0.83 | Maxes |
| BC | 0.63 | 1.00 | 0.94 | 0.80 | |
| PBMC | 0.45 | 0.79 | 1.00 | 0.79 | |
| Pheno Age | 0.66 | 0.66 | 0.55 | 1.00 | |
| | Mins | | | | |

| DNAmTL | Saliva | BC | PBMC | Pheno Age | |
|---|---|---|---|---|---|
| Saliva | 1.00 | 0.31 | 0.28 | 0.07 | Maxes |
| BC | -0.17 | 1.00 | 0.79 | -0.14 | |
| PBMC | -0.22 | 0.56 | 1.00 | -0.01 | |
| Pheno Age | -0.32 | -0.50 | -0.49 | 1.00 | |

| epiTOC | Saliva | BC | PBMC | Pheno Age | |
|---|---|---|---|---|---|
| Saliva | 1.00 | 0.46 | 0.34 | 0.24 | Maxes |
| BC | -0.17 | 1.00 | 0.86 | 0.23 | |
| PBMC | -0.22 | 0.68 | 1.00 | 0.36 | |
| Pheno Age | -0.14 | -0.09 | -0.15 | 1.00 | |

| epiTOC2 | Saliva | BC | PBMC | Pheno Age |
|---|---|---|---|---|
| Saliva | 1.00 | 0.38 | 0.24 | 0.23 |
| BC | -0.23 | 1.00 | 0.89 | 0.28 |
| PBMC | -0.25 | 0.68 | 1.00 | 0.31 |
| Pheno Age | -0.06 | -0.06 | -0.17 | 1.00 |

Mins (below diagonal); Maxes (above diagonal)

| Pan-Mam2 | Saliva | BC | PBMC | Pheno Age |
|---|---|---|---|---|
| Saliva | 1.00 | 0.49 | 0.40 | 0.55 |
| BC | 0.13 | 1.00 | 0.78 | 0.69 |
| PBMC | -0.01 | 0.50 | 1.00 | 0.44 |
| Pheno Age | 0.25 | 0.46 | 0.10 | 1.00 |

Mins (below diagonal); Maxes (above diagonal)

| Pan-Mam3 | Saliva | BC | PBMC | Pheno Age |
|---|---|---|---|---|
| Saliva | 1.00 | 0.38 | 0.43 | 0.58 |
| BC | 0.01 | 1.00 | 0.73 | 0.70 |
| PBMC | 0.05 | 0.55 | 1.00 | 0.50 |
| Pheno Age | 0.18 | 0.39 | 0.13 | 1.00 |

Mins (below diagonal); Maxes (above diagonal)

| PedBE | Saliva | BC | PBMC | Pheno Age |
|---|---|---|---|---|
| Saliva | 1.00 | 0.51 | 0.45 | 0.60 |
| BC | 0.06 | 1.00 | 0.66 | 0.39 |
| PBMC | 0.01 | 0.30 | 1.00 | 0.59 |
| Pheno Age | 0.26 | -0.03 | 0.30 | 1.00 |

Mins (below diagonal); Maxes (above diagonal)

| ZhangY | Saliva | BC | PBMC | Pheno Age |
|---|---|---|---|---|
| Saliva | 1.00 | 0.50 | 0.27 | 0.39 |
| BC | 0.09 | 1.00 | 0.72 | 0.48 |
| PBMC | -0.20 | 0.42 | 1.00 | 0.45 |
| Pheno Age | 0.03 | 0.17 | 0.11 | 1.00 |

Mins (below diagonal); Maxes (above diagonal)

BC = buffy coat; PBMC = peripheral blood mononuclear cells; Pheno = Phenotypic.

**Table S3** - Linear Mixed-effects Model Fit of Fixed Effects, DNA Methylation Clocks by Sample Medium.

| DNA Methylation Clock | *B* | *SE* | df | *p*-value | Between Family SD | Within Family SD |
|---|---|---|---|---|---|---|
| **9. DNAmTL** | | | | | | |
| Intercept [BC=ref] | 7.354 | 0.021 | 212 | <0.001 | 0.085 | 0.166 |
| PBMC | -0.007 | 0.025 | 212 | 0.789 | | |
| Saliva | -0.711 | 0.025 | 212 | <0.001 | | |
| Chronological Age | -0.016 | 0.004 | 212 | <0.001 | | |
| **10. epiTOC** | | | | | | |
| Intercept [BC=ref] | 0.059 | 0.001 | 212 | <0.001 | 0.004 | 0.006 |
| PBMC | <-0.001 | 0.001 | 212 | 0.922 | | |
| Saliva | 0.019 | 0.001 | 212 | <0.001 | | |
| Chronological Age | <0.001 | <0.001 | 212 | 0.125 | | |
| **11. epiTOC2** | | | | | | |
| Intercept [BC=ref] | 1497.2 | 44.8 | 212 | <0.001 | 192.288 | 335.528 |
| PBMC | -6.7 | 49.7 | 212 | 0.892 | | |
| Saliva | 817.2 | 51.2 | 212 | <0.001 | | |
| Chronological Age | 15.4 | 8.8 | 212 | 0.083 | | |
| **12. Pan-Mammalian2** | | | | | | |
| Intercept [BC=ref] | 45.912 | 0.304 | 212 | <0.001 | 1.242 | 2.340 |
| PBMC | -0.034 | 0.347 | 212 | 0.922 | | |
| Saliva | 11.833 | 0.357 | 212 | <0.001 | | |
| Chronological Age | 0.369 | 0.059 | 212 | <0.001 | | |
| **13. Pan-Mammalian3** | | | | | | |
| Intercept [BC=ref] | 50.159 | 0.297 | 212 | <0.001 | 1.102 | 2.393 |
| PBMC | 0.023 | 0.355 | 212 | 0.948 | | |
| Saliva | 5.985 | 0.365 | 212 | <0.001 | | |
| Chronological Age | 0.311 | 0.056 | 212 | <0.001 | | |
| **14. PedBE** | | | | | | |
| Intercept [BC=ref] | 7.698 | 0.154 | 212 | <0.001 | 0.459 | 1.328 |
| PBMC | -0.007 | 0.197 | 212 | 0.973 | | |
| Saliva | 10.680 | 0.203 | 212 | <0.001 | | |
| Chronological Age | 0.137 | 0.028 | 212 | <0.001 | | |
| **15. ZhangY** | | | | | | |
| Intercept [BC=ref] | -1.914 | 0.027 | 212 | <0.001 | 0.141 | 0.169 |
| PBMC | 0.025 | 0.025 | 212 | 0.315 | | |
| Saliva | 1.061 | 0.026 | 212 | <0.001 | | |
| Chronological Age | 0.020 | 0.006 | 212 | <0.001 | | |

*Notes.* BC = buffy coat; PBMC = peripheral blood mononuclear cells; SD = standard deviation.

**Table S4** - Linear Mixed-effects Results DNA Methylation Clocks and Age Accelerations by Sample Medium without Chronological Age.

| | DNAm Age/Clocks | | | | | | DNAm Age Acceleration | | | | | |
|---|---|---|---|---|---|---|---|---|---|---|---|---|
| **DNA Methylation Clock** | ***B*** | ***SE*** | **df** | **p-value** | **Between Family SD** | **Within Family SD** | ***B*** | ***SE*** | **df** | **p-value** | **Between Family SD** | **Within Family SD** |
| **1. Cortical Clock** | | | | | | | | | | | | |
| Intercept [BC=ref] | 38.136 | 0.457 | 213 | <0.001 | 2.465 | 2.763 | -3.833 | 0.367 | 213 | <0.001 | 1.635 | 2.683 |
| PBMC | -0.057 | 0.410 | 213 | 0.890 | | | -0.057 | 0.398 | 213 | 0.886 | | |
| Saliva | 12.371 | 0.422 | 213 | <0.001 | | | 12.344 | 0.410 | 213 | <0.001 | | |
| **2. DunedinPACE** | | | | | | | | | | | | |
| Intercept [BC=ref] | 1.304 | 0.011 | 213 | <0.001 | 0.056 | 0.070 | 0.000 | 0.011 | 213 | 0.973 | 0.056 | 0.070 |
| PBMC | 0.003 | 0.010 | 213 | 0.786 | | | 0.003 | 0.010 | 213 | 0.785 | | |
| Saliva | -0.007 | 0.011 | 213 | 0.497 | | | -0.007 | 0.011 | 213 | 0.492 | | |
| **3. Hannum** | | | | | | | | | | | | |
| Intercept [BC=ref] | 36.167 | 0.499 | 213 | <0.001 | 2.832 | 2.766 | -3.942 | 0.363 | 213 | <0.001 | 1.582 | 2.692 |
| PBMC | 0.136 | 0.410 | 213 | 0.741 | | | 0.136 | 0.399 | 213 | 0.734 | | |
| Saliva | 12.557 | 0.423 | 213 | <0.001 | | | 12.514 | 0.411 | 213 | <0.001 | | |
| **4. Horvath** | | | | | | | | | | | | |
| Intercept [BC=ref] | 22.002 | 0.627 | 213 | <0.001 | 3.541 | 3.514 | -2.476 | 0.468 | 213 | <0.001 | 2.088 | 3.416 |
| PBMC | 0.386 | 0.521 | 213 | 0.459 | | | 0.386 | 0.506 | 213 | 0.446 | | |
| Saliva | 7.470 | 0.537 | 213 | <0.001 | | | 7.424 | 0.522 | 213 | <0.001 | | |
| **5. Horvath2** | | | | | | | | | | | | |
| Intercept [BC=ref] | 18.604 | 0.584 | 213 | <0.001 | 3.405 | 3.064 | -5.154 | 0.393 | 213 | <0.001 | 1.675 | 2.962 |
| PBMC | 0.085 | 0.454 | 213 | 0.852 | | | 0.085 | 0.439 | 213 | 0.847 | | |
| Saliva | 16.455 | 0.468 | 213 | <0.001 | | | 16.412 | 0.452 | 213 | <0.001 | | |
| **6. PCGrimAge** | | | | | | | | | | | | |

| | | | | | | | | | | | | |
|---|---|---|---|---|---|---|---|---|---|---|---|---|
| Intercept [BC=ref] | 48.399 | 0.534 | 213 | <0.001 | 3.802 | 1.428 | -2.764 | 0.287 | 213 | <0.001 | 1.810 | 1.174 |
| PBMC | 0.135 | 0.212 | 213 | 0.524 | | | 0.135 | 0.174 | 213 | 0.438 | | |
| Saliva | 9.158 | 0.218 | 213 | <0.001 | | | 9.107 | 0.179 | 213 | <0.001 | | |
| **7. PhenoAge** | | | | | | | | | | | | |
| Intercept [BC=ref] | 20.365 | 0.842 | 213 | <0.001 | 4.115 | 5.710 | -6.273 | 0.705 | 213 | <0.001 | 2.710 | 5.594 |
| PBMC | 0.269 | 0.847 | 213 | 0.751 | | | 0.269 | 0.829 | 213 | 0.746 | | |
| Saliva | 19.791 | 0.872 | 213 | <0.001 | | | 19.743 | 0.854 | 213 | <0.001 | | |
| **8. ZhangQ** | | | | | | | | | | | | |
| Intercept [BC=ref] | 26.423 | 0.619 | 213 | <0.001 | 3.887 | 2.591 | -1.235 | 0.344 | 213 | <0.001 | 1.617 | 2.412 |
| PBMC | 0.217 | 0.384 | 213 | 0.573 | | | 0.217 | 0.358 | 213 | 0.545 | | |
| Saliva | 3.832 | 0.396 | 213 | <0.001 | | | 3.770 | 0.368 | 213 | <0.001 | | |
| **9. DNAmTL** | | | | | | | | | | | | |
| Intercept [BC=ref] | 7.351 | 0.023 | 213 | <0.001 | 0.098 | 0.168 | 0.221 | 0.021 | 213 | <0.001 | 0.084 | 0.166 |
| PBMC | -0.006 | 0.025 | 213 | 0.791 | | | -0.007 | 0.025 | 213 | 0.789 | | |
| Saliva | -0.712 | 0.026 | 213 | <0.001 | | | -0.711 | 0.025 | 213 | <0.001 | | |
| **10. epiTOC** | | | | | | | | | | | | |
| Intercept [BC=ref] | 0.059 | 0.001 | 213 | <0.001 | 0.004 | 0.006 | -0.006 | 0.001 | 213 | <0.001 | 0.004 | 0.006 |
| PBMC | <-0.001 | 0.001 | 213 | 0.922 | | | <-0.001 | 0.001 | 213 | 0.922 | | |
| Saliva | 0.019 | 0.001 | 213 | <0.001 | | | 0.019 | 0.001 | 213 | <0.001 | | |
| **11. epiTOC2** | | | | | | | | | | | | |
| Intercept [BC=ref] | 1499.7 | 45.1 | 213 | <0.001 | 195.4 | 336.3 | -253.1 | 44.5 | 213 | 0.000 | 189.4 | 335.5 |
| PBMC | -6.7 | 49.9 | 213 | 0.893 | | | -6.7 | 49.7 | 213 | 0.892 | | |
| Saliva | 817.9 | 51.4 | 213 | <0.001 | | | 817.3 | 51.2 | 213 | <0.001 | | |
| **12. Pan-Mammalian2** | | | | | | | | | | | | |
| Intercept [BC=ref] | 45.975 | 0.360 | 213 | <0.001 | 1.811 | 2.369 | -3.688 | 0.302 | 213 | <0.001 | 1.223 | 2.340 |
| PBMC | -0.034 | 0.351 | 213 | 0.923 | | | -0.034 | 0.347 | 213 | 0.922 | | |

| | | | | | | | | | | | | |
|---|---|---|---|---|---|---|---|---|---|---|---|---|
| Saliva | 11.855 | 0.362 | 213 | <0.001 | | | 11.832 | 0.357 | 213 | <0.001 | | |
| **13. Pan-Mammalian3** | | | | | | | | | | | | |
| Intercept [BC=ref] | 50.215 | 0.337 | 213 | <0.001 | 1.542 | 2.421 | -1.860 | 0.295 | 213 | <0.001 | 1.082 | 2.393 |
| PBMC | 0.023 | 0.359 | 213 | 0.948 | | | 0.023 | 0.355 | 213 | 0.948 | | |
| Saliva | 6.002 | 0.370 | 213 | <0.001 | | | 5.985 | 0.365 | 213 | <0.001 | | |
| **14. PedBE** | | | | | | | | | | | | |
| Intercept [BC=ref] | 7.712 | 0.169 | 213 | <0.001 | 0.657 | 1.340 | -3.320 | 0.154 | 213 | <0.001 | 0.448 | 1.328 |
| PBMC | -0.007 | 0.199 | 213 | 0.974 | | | -0.007 | 0.197 | 213 | 0.973 | | |
| Saliva | 10.687 | 0.204 | 213 | <0.001 | | | 10.680 | 0.203 | 213 | <0.001 | | |
| **15. ZhangY** | | | | | | | | | | | | |
| Intercept [BC=ref] | -1.909 | 0.028 | 213 | <0.001 | 0.154 | 0.171 | -0.331 | 0.027 | 213 | <0.001 | 0.139 | 0.169 |
| PBMC | 0.025 | 0.025 | 213 | 0.996 | | | 0.025 | 0.025 | 213 | 0.316 | | |
| Saliva | 1.062 | 0.026 | 213 | <0.001 | | | 1.061 | 0.026 | 213 | <0.001 | | |

*Note*. DNAm=DNA methylation; BC = buffy coat; PBMC = peripheral blood mononuclear cells; SD = standard deviation.

**Table S5.** Twin Spearman correlations for age acceleration residuals, rate of aging, and mortality risk measures.

| | | Spearman correlations (random twin assignment, 100 iterations) | | | | | | | | Cross-tissue correlations | | | |
|---|---|---|---|---|---|---|---|---|---|---|---|---|---|
| **DNA Methylation Clock** | **Tissue** | **Mean rMZ** | **SD rMZ** | **95% range of rMZ** | **% p<0.05** | **Mean rDZ** | **SD rDZ** | **95% range of rDZ** | **% p<0.05** | $\overline{r}_{s,MZ}$ | $SD_{MZ}$ | $\overline{r}_{s,DZ}$ | $SD_{DZ}$ |
| 1. CorticalClock | Saliva | 0.37 | 0.06 | (0.26, 0.49) | 1% | **0.72** | **0.05** | **(0.65, 0.83)** | **100%** | | | | |
| | BC | **0.66** | **0.03** | **(0.60-0.72)** | **100%** | 0.09 | 0.06 | (0.01, 0.23) | 0% | | | | |
| | PBMC | 0.27 | 0.05 | (0.19-0.38) | 0% | 0.36 | 0.06 | (0.26-0.49) | 1% | 0.43 | 0.20 | 0.39 | 0.32 |
| 2. DunedinPACE* | Saliva | 0.53 | 0.05 | (0.45, 0.64) | 38% | 0.42 | 0.09 | (0.28, 0.60) | 4% | | | | |
| | BC | **0.70** | **0.04** | **(0.64, 0.79** | **100%** | 0.24 | 0.09 | (0.11, 0.50) | 3% | | | | |
| | PBMC | **0.59** | **0.04** | **(0.53, 0.67)** | **100%** | 0.10 | 0.09 | (-0.03, 0.29) | 0% | 0.61 | 0.09 | 0.25 | 0.16 |
| 3. Hannum | Saliva | 0.11 | 0.08 | (-0.04, 0.29) | 0% | 0.57 | 0.05 | (0.50, 0.66) | 37% | | | | |
| | BC | **0.57** | **0.05** | **(0.49-0.67)** | **99%** | 0.36 | 0.08 | (0.24-0.51) | 2% | | | | |
| | PBMC | 0.37 | 0.05 | (0.29-0.50) | 5% | **0.70** | **0.05** | **(0.60, 0.82)** | **100%** | 0.35 | 0.23 | 0.54 | 0.17 |
| 4. Horvath | Saliva | 0.19 | 0.08 | (0.06, 0.35) | 0% | 0.42 | 0.06 | (0.31, 0.55) | 0% | | | | |
| | BC | **0.64** | **0.04** | **(0.58-0.75)** | **100%** | 0.31 | 0.05 | (0.23-0.42) | 0% | | | | |
| | PBMC | 0.28 | 0.04 | (0.20, 0.38) | 0% | **0.65** | **0.05** | **(0.57-0.75)** | **99%** | 0.37 | 0.24 | 0.46 | 0.17 |
| 5. Horvath2 | Saliva | 0.28 | 0.07 | (0.14, 0.42) | 0% | 0.47 | 0.08 | (0.31, 0.66) | 10% | | | | |
| | BC | **0.71** | **0.04** | **(0.65-0.78)** | **100%** | 0.50 | 0.06 | (0.41-0.64) | 21% | | | | |
| | PBMC | **0.53** | **0.04** | **(0.47-0.61)** | **93%** | **0.67** | **0.04** | **(0.57-0.75)** | **100%** | 0.51 | 0.22 | 0.55 | 0.11 |
| 6. PCGrimAge | Saliva | **0.86** | **0.04** | **(0.78, 0.93)** | **100%** | **0.59** | **0.06** | **(0.51, 0.73)** | **45%** | | | | |
| | BC | **0.80** | **0.02** | **(0.76-0.83)** | **100%** | 0.50 | 0.05 | (0.41-0.59) | 13% | | | | |
| | PBMC | **0.90** | **0.02** | **(0.86-0.93)** | **100%** | 0.35 | 0.05 | (0.26-0.47) | 0% | 0.85 | 0.05 | 0.48 | 0.12 |

| DNA Methylation Clock | Tissue | Spearman correlations (random twin assignment, 100 iterations) | | | | | | | | Cross-tissue correlations | | | |
|---|---|---|---|---|---|---|---|---|---|---|---|---|---|
| | | Mean rMZ | SD rMZ | 95% range of rMZ | % p<0.05 | Mean rDZ | SD rDZ | 95% range of rDZ | % p<0.05 | $\overline{r}_{s,MZ}$ | $SD_{MZ}$ | $\overline{r}_{s,DZ}$ | $SD_{DZ}$ |
| 7. PhenoAge | Saliva | 0.39 | 0.08 | (0.22, 0.58) | 4% | 0.18 | 0.05 | (0.10, 0.25) | 0% | | | | |
| | BC | **0.55** | **0.05** | **(0.46-0.66)** | **100%** | 0.49 | 0.05 | (0.39-0.57) | 13% | | | | |
| | PBMC | 0.21 | 0.07 | (0.11-0.37) | 1% | 0.41 | 0.05 | (0.32-0.51) | 2% | 0.38 | 0.17 | 0.36 | 0.16 |
| 8. ZhangQ | Saliva | 0.22 | 0.11 | (0.05, 0.41) | 2% | 0.38 | 0.09 | (0.26, 0.60) | 4% | | | | |
| | BC | **0.76** | **0.03** | **(0.71, 0.82)** | **100%** | **0.69** | **0.04** | **(0.61, 0.78)** | **100%** | | | | |
| | PBMC | **0.57** | **0.05** | **(0.49-0.67)** | **99%** | **0.61** | **0.06** | **(0.50-0.73)** | **91%** | 0.52 | 0.27 | 0.56 | 0.16 |
| 9. DNAmTL* | Saliva | -0.01 | 0.07 | (-0.13, 0.18) | 0% | 0.22 | 0.08 | (0.10, 0.38) | 0% | | | | |
| | BC | 0.42 | 0.04 | (0.34-0.50) | 8% | 0.22 | 0.07 | (0.12-0.38) | 0% | | | | |
| | PBMC | 0.42 | 0.05 | (0.34-0.52) | 17% | 0.17 | 0.07 | (0.05-0.34) | 0% | 0.28 | 0.25 | 0.20 | 0.03 |
| 10. epiTOC* | Saliva | 0.47 | 0.05 | (0.38, 0.56) | 7% | 0.05 | 0.10 | (-0.09, 0.31) | 0% | | | | |
| | BC | **0.71** | **0.03** | **(0.66-0.78)** | **100%** | 0.54 | 0.05 | (0.45-0.66) | 38% | | | | |
| | PBMC | **0.60** | **0.05** | **(0.53, 0.71)** | **99%** | 0.28 | 0.07 | (0.17, 0.45) | 0% | 0.59 | 0.12 | 0.29 | 0.25 |
| 11. epiTOC2* | Saliva | 0.51 | 0.03 | (0.46, 0.56) | 23% | 0.01 | 0.09 | (-0.15, 0.21) | 0% | | | | |
| | BC | **0.65** | **0.04** | **(0.57-0.71)** | **100%** | 0.52 | 0.07 | (0.42, 0.71) | 33% | | | | |
| | PBMC | **0.60** | **0.04** | **(0.53-0.71)** | **100%** | 0.24 | 0.08 | (0.11, 0.48) | 0% | 0.59 | 0.07 | 0.26 | 0.26 |
| 12. Pan-Mammalian Clock2 | Saliva | -0.04 | 0.07 | (-0.15, 0.12) | 0% | 0.32 | 0.11 | (0.18, 0.59) | 3% | | | | |
| | BC | **0.61** | **0.05** | **(0.54-0.70)** | **100%** | 0.46 | 0.08 | (0.34-0.69) | 9% | | | | |
| | PBMC | **0.71** | **0.04** | **(0.66-0.80)** | **100%** | 0.32 | 0.06 | (0.22-0.46) | 0% | 0.43 | 0.41 | 0.37 | 0.08 |

| DNA Methylation Clock | Tissue | Spearman correlations (random twin assignment, 100 iterations) | | | | | | | | Cross-tissue correlations | | | |
|---|---|---|---|---|---|---|---|---|---|---|---|---|---|
| | | Mean rMZ | SD rMZ | 95% range of rMZ | % p<0.05 | Mean rDZ | SD rDZ | 95% range of rDZ | % p<0.05 | $\overline{r}_{s,MZ}$ | $SD_{MZ}$ | $\overline{r}_{s,DZ}$ | $SD_{DZ}$ |
| 13. Pan-Mammalian Clock3 | Saliva | 0.17 | 0.04 | (0.10, 0.24) | 0% | -0.05 | 0.10 | (-0.20, 0.17) | 0% | | | | |
| | BC | **0.64** | **0.04** | **(0.57-0.74)** | **100%** | **0.64** | **0.04** | **(0.57-0.71)** | **100%** | | | | |
| | PBMC | **0.54** | **0.05** | **(0.46-0.63)** | **92%** | 0.27 | 0.07 | (0.17-0.41) | 0% | 0.45 | 0.25 | 0.29 | 0.35 |
| 14. PedBE | Saliva | 0.22 | 0.07 | (0.10, 0.36) | 0% | -0.11 | 0.07 | (-0.23, 0.03) | 0% | | | | |
| | BC | **0.67** | **0.05** | **(0.59-0.79)** | **100%** | -0.02 | 0.08 | (-0.14, 0.16) | 0% | | | | |
| | PBMC | 0.36 | 0.05 | (0.28-0.47) | 3% | **0.59** | **0.05** | **(0.52-0.68)** | **90%** | 0.42 | 0.23 | 0.15 | 0.38 |
| 15. ZhangY* | Saliva | **0.64** | **0.04** | **(0.55, 0.72)** | **99%** | 0.22 | 0.09 | (0.06, 0.45) | 0% | | | | |
| | BC | 0.45 | 0.05 | (0.37-0.53) | 31% | 0.52 | 0.06 | (0.43-0.66) | 27% | | | | |
| | PBMC | 0.41 | 0.05 | (0.30-0.52) | 13% | **0.77** | **0.05** | **(0.67-0.86)** | **100%** | 0.50 | 0.12 | 0.50 | 0.28 |

*Indicates absolute DNAm measures used in correlations; All others are age acceleration residuals.
*Note*. DNAm=DNA methylation; BC= Buffy Coat; PBMC= peripheral blood mononuclear cells. Bolded values indicate 90-100% of the randominized tests were $p<0.05$. $N_{MZ}$=18 pairs for BC and PBMC and $N_{MZ}$=14 pairs for Saliva; $N_{DZ}$=14 pairs for BC and PBMC and $N_{DZ}$=12 pairs for Saliva.

### References


Lu, A. T., Fei, Z., Haghani, A., Robeck, T. R., Zoller, J. A., Li, C. Z., Lowe, R., Yan, Q., Zhang, J., Vu, H., Ablaeva, J., Acosta-Rodriguez, V. A., Adams, D. M., Almunia, J., Aloysius, A., Ardehali, R., Arneson, A., Baker, C. S., Banks, G., Belov, K., Bennett, N. C., Black, P., Blumstein, D. T., Bors, E. K., Breeze, C. E., Brooke, R. T., Brown, J. L., Carter, G. G., Caulton, A., Cavin, J. M., Chakrabarti, L., Chatzistamou, I., Chen, H., Cheng, K., Chiavellini, P., Choi, O. W., Clarke, S. M., Cooper, L. N., Cossette, M. L., Day, J., DeYoung, J., DiRocco, S., Dold, C., Ehmke, E. E., Emmons, C. K., Emmrich, S., Erbay, E., Erlacher-Reid, C., Faulkes, C. G., Ferguson, S. H., Finno, C. J., Flower, J. E., Gaillard, J. M., Garde, E., Gerber, L., Gladyshev, V. N., Gorbunova, V., Goya, R. G., Grant, M. J., Green, C. B., Hales, E. N., Hanson, M. B., Hart, D. W., Haulena, M., Herrick, K., Hogan, A. N., Hogg, C. J., Hore, T. A., Huang, T., Izpisua Belmonte, J. C., Jasinska, A. J., Jones, G., Jourdain, E., Kashpur, O., Katcher, H., Katsumata, E., Kaza, V., Kiaris, H., Kobor, M. S., Kordowitzki, P., Koski, W. R., Krutzen, M., Kwon, S. B., Larison, B., Lee, S. G., Lehmann, M., Lemaitre, J. F., Levine, A. J., Li, C., Li, X., Lim, A. R., Lin, D. T. S., Lindemann, D. M., Little, T. J., Macoretta, N., Maddox, D., Matkin, C. O., Mattison, J. A., McClure, M., Mergl, J., Meudt, J. J., Montano, G. A., Mozhui, K., Munshi-South, J., Naderi, A., Nagy, M., Narayan, P., Nathanielsz, P. W., Nguyen, N. B., Niehrs, C., O'Brien, J. K., O'Tierney Ginn, P., Odom, D. T., Ophir, A. G., Osborn, S., Ostrander, E. A., Parsons, K. M., Paul, K. C., Pellegrini, M., Peters, K. J., Pedersen, A. B., Petersen, J. L., Pietersen, D. W., Pinho, G. M., Plassais, J., Poganik, J. R., Prado, N. A., Reddy, P., Rey, B., Ritz, B. R., Robbins, J., Rodriguez, M., Russell, J., Rydkina, E., Sailer, L. L., Salmon, A. B., Sanghavi, A., Schachtschneider, K. M., Schmitt, D., Schmitt, T., Schomacher, L., Schook, L. B., Sears, K. E., Seifert, A. W., Seluanov, A., Shafer, A. B. A., Shanmuganayagam, D., Shindyapina, A. V., Simmons, M., Singh, K., Sinha, I., Slone, J., Snell, R. G., Soltanmaohammadi, E., Spangler, M. L., Spriggs, M. C., Staggs, L., Stedman, N., Steinman, K. J., Stewart, D. T., Sugrue, V. J., Szladovits, B., Takahashi, J. S., Takasugi, M., Teeling, E. C., Thompson, M. J., Van Bonn, B., Vernes, S. C., Villar, D., Vinters, H. V., Wallingford, M. C., Wang, N., Wayne, R. K., Wilkinson, G. S., Williams, C. K., Williams, R. W., Yang, X. W., Yao, M., Young, B. G., Zhang, B., Zhang, Z., Zhao, P., Zhao, Y., Zhou, W., Zimmermann, J., Ernst, J., Raj, K., & Horvath, S. (2023). Author Correction: Universal DNA methylation age across mammalian tissues. *Nat Aging, 3*(11), 1462. doi:10.1038/s43587-023-00499-7

Lu, A. T., Seeboth, A., Tsai, P. C., Sun, D., Quach, A., Reiner, A. P., Kooperberg, C., Ferrucci, L., Hou, L., Baccarelli, A. A., Li, Y., Harris, S. E., Corley, J., Taylor, A., Deary, I. J., Stewart, J. D., Whitsel, E. A., Assimes, T. L., Chen, W., Li, S., Mangino, M., Bell, J. T., Wilson, J. G., Aviv, A., Marioni, R. E., Raj, K., & Horvath, S. (2019). DNA methylation-based estimator of telomere length. *Aging (Albany NY), 11*(16), 5895-5923. doi:10.18632/aging.102173

McEwen, L. M., O'Donnell, K. J., McGill, M. G., Edgar, R. D., Jones, M. J., MacIsaac, J. L., Lin, D. T. S., Ramadori, K., Morin, A., Gladish, N., Garg, E., Unternaehrer, E., Pokhvisneva, I., Karnani, N., Kee, M. Z. L., Klengel, T., Adler, N. E., Barr, R. G., Letourneau, N., Giesbrecht, G. F., Reynolds, J. N., Czamara, D., Armstrong, J. M., Essex, M. J., de Weerth, C., Beijers, R., Tollenaar, M. S., Bradley, B., Jovanovic, T., Ressler, K. J., Steiner, M., Entringer, S., Wadhwa, P. D., Buss, C., Bush, N. R., Binder, E. B., Boyce, W. T., Meaney, M. J., Horvath, S., & Kobor, M. S. (2020). The PedBE clock accurately estimates DNA methylation age in pediatric buccal cells. *Proc Natl Acad Sci U S A, 117*(38), 23329-23335. doi:10.1073/pnas.1820843116

Teschendorff, A. E. (2020). A comparison of epigenetic mitotic-like clocks for cancer risk prediction. *Genome Med, 12*(1), 56. doi:10.1186/s13073-020-00752-3
Yang, Z., Wong, A., Kuh, D., Paul, D. S., Rakyan, V. K., Leslie, R. D., Zheng, S. C., Widschwendter, M., Beck, S., & Teschendorff, A. E. (2016). Correlation of an epigenetic mitotic clock with cancer risk. *Genome Biol, 17*(1), 205. doi:10.1186/s13059-016-1064-3
Zhang, Y., Wilson, R., Heiss, J., Breitling, L. P., Saum, K. U., Schottker, B., Holleczek, B., Waldenberger, M., Peters, A., & Brenner, H. (2017). DNA methylation signatures in peripheral blood strongly predict all-cause mortality. *Nat Commun, 8*(1), 14617. doi:10.1038/ncomms14617
